# Structural elucidation of the hexameric MmpS4-MmpL4 complex from *Mycobacterium tuberculosis*

**DOI:** 10.64898/2026.01.07.698164

**Authors:** Jennifer C. Earp, Nicolas P. Lichti, Alisa A. Garaeva, Virginia Meikle, Michael Niederweis, Markus A. Seeger

## Abstract

*Mycobacterium tuberculosis* contains thirteen Mycobacterial membrane protein Large (MmpL) transporters, which belong to the family of secondary active RND transporters. MmpL4 and MmpL5, together with their operon partners MmpS4 and MmpS5, export the mycobacterial siderophore mycobactin and the last resort TB drug bedaquiline. Recently, we determined a structure of the MmpL4 monomer in complex with desferrated mycobactin, which lacked a functionally essential coiled-coil domain predicted to extend far into the periplasm. Here, we present a cryo-EM structure of the hexameric (MmpS4)_3_-(MmpL4)_3_ complex, which was enabled by rational disulfide cross-links based on AlphaFold predictions. We observed density for the coiled-coil domain, which protrudes into the periplasmic space at an angle of around 60° relative to the symmetry axis of the MmpL4 trimer. In the context of the hexameric complex, MmpL4’s conformation differs strikingly from the one observed for monomeric MmpL4, which includes formation of a large cavity in the periplasmic domain and rearrangements of conserved proton coupling residues at the transmembrane domain. Two of the cross-linked variants were found to be functionally defective, indicating that the hexameric complex undergoes substantial conformational changes to accomplish mycobactin and drug transport.

## INTRODUCTION

*Mycobacterium tuberculosis* encodes 13 resistance-nodulation-cell division (RND) transporters, referred to as the Mycobacterial membrane protein Large (MmpL) family [1]. Multiple MmpL transporters are of interest as novel targets for anti-TB drugs, including the only essential member of the MmpL family, the trehalose monomycolate (TMM) exporter MmpL3 [2–7], and the two siderophore (mycobactin) exporters MmpL4 and MmpL5, which are essential under low iron conditions and strongly contribute to virulence in mice [8, 9]. Importantly, MmpL4 and in particular MmpL5 also confer resistance to bedaquiline, an antibiotic crucial for the treatment of multidrug resistant *M. tuberculosis* [10–13].

Like other RND transporters, MmpL proteins are proton-driven antiporters composed of a transmembrane domain and a periplasmic domain [2, 3]. The transmembrane domain consists of two pseudosymmetric six-helix bundles and contains conserved residues that are crucial for proton translocation. In case of MmpL3, MmpL4 and MmpL5, two DY-pairs located at the center of the transmembrane domain have been suggested to enable proton coupling [14–16]. The periplasmic domain is made up of two domains, referred to as the N-terminal (PN) and and C-terminal (PC) porter domain, which adopt different conformations in MmpL3 and MmpL4 [4, 15]. However, the mechanistic basis of how conformational changes in the periplasmic domain are coupled to protonation of the DY-pairs in the transmembrane domain remains elusive.

MmpL proteins are divided into two clusters. Cluster I MmpL transporters (including MmpL4 and MmpL5) contain an additional coiled-coil domain (CCD), a structural element extending into the periplasmic space. Cluster II MmpL transporters (including MmpL3) lack this domain. The CCD was first described based on the AlphaFold prediction of MmpL4, and was proposed to be involved in multimerization of MmpL proteins [15]. A subset of cluster I MmpL transporters (including MmpL4 and MmpL5) are associated with a periplasmic accessory protein called Mycobacterial membrane protein small (MmpS) which is encoded in the same operon.

Structures of cluster I MmpL transporters were determined only recently. *M. smegmatis* MSMEG_1382 was the first published structure of a cluster I MmpL transporter, but the CCD was not resolved due to the flexibility of the domain [17]. The second published cryo-EM structure was that of monomeric *M. tuberculosis* MmpL4 in complex with its substrate mycobactin, determined by our lab [15]. To obtain this structure the CCD was truncated, because full-length MmpL4 was not amenable for structure determination [15]. We also demonstrated that the CCDs of MmpL4 and MmpL5 are essential for mycobactin and bedaquiline export in *M. tuberculosis* [15]. Biochemical studies suggested that the CCD is responsible for trimer formation of *M. smegmatis* MmpL10 transporter (MSMEG_0410) [18], while TIRF microscopy showed that MmpS5 is required for MmpL5 trimer formation [19]. In agreement with these findings, recent structures of hexameric *M. tuberculosis* MmpS5-MmpL5 and *M. smegmatis* MSMEG_1381-MSMEG_1382 complexes confirmed that both the CCD and MmpS protein contribute to MmpL trimerization [16, 20]. Even though the CCD and MmpS contribute to MmpL trimerization, neither are strictly required to obtain a stable multimeric assembly of the MmpL complex. This was demonstrated by a structure of hexameric *M. tuberculosis* MmpS5-MmpL5 wherein the CCD was genetically deleted [21] and purification of trimers of full-length MmpL5 in the absence of MmpS5 [16].

MmpL4, MmpL5 and MSMEG_1382 co-purify with abundant acyl-carrier proteins (ACPs) from the respective expression host, namely *E. coli* ACP with MmpL4 [15] or AcpM with MmpL5 [16] and MSMEG_1382 [22]. Biochemical analyses showed that MmpL4 also interacts with MbtL (Rv1344), an ACP involved in mycobactin biosynthesis, suggesting that mycobactin biosynthesis and export are coupled processes [15].

In this work, we utilized the AlphaFold2 model of the (MmpS4)_3_-(MmpL4)_3_ complex to design disulfide bonds between MmpS4 and MmpL4. These engineered disulfide cross-links stabilized the MmpS4-MmpL4 interface, thereby enabling cryo-EM structure determination of hexameric MmpS4-MmpL4. The structure revealed a binding cavity in the periplasmic domain of MmpL4.

## RESULTS

### Prediction of a hexameric MmpS4-MmpL4 complex using AlphaFold2

In order to purify MmpL4 from *E. coli*, *mmpL4* had to be expressed together with its operon partner *mmpS4* [15]. When full-length MmpL4 (including its CCD) was purified via a C-terminal GFP-tag, the sample predominantly eluted as a broad peak from the size exclusion chromatography (SEC) column, corresponding to MmpL4 multimers of varying stoichiometries [15]. A later-eluting peak contained the MmpL4 monomer and suffered from protein degradation. SDS-PAGE analysis and mass-spectrometry analyses of the early-eluting peak revealed that MmpS4 was co-purified, as well as the acyl-carrier protein ACP of the expression host *E. coli.* However, when attempting to determine a cryo-EM structure of the multimeric MmpS4-MmpL4 complex, the sample quality was poor, only permitting reconstruction of a 6.3 Å map of the MmpL4 monomer in complex with ACP [15]. Hence, in our hands, the detergent-purified MmpS4-MmpL4 complex was not stable enough for high resolution structural analysis.

Therefore, we used AlphaFold2 to predict the multimeric complex including three MmpL4 and three MmpS4 chains [23]. AlphaFold2 generated a model with high values for the predicted local distance difference test (pLDDT) score and importantly also for the predicted alignment error (PAE) over the entire polypeptide chains of MmpL4 and MmpS4 indicating a high confidence in the predicted interactions (Figure 1A, Figure 1 — figure supplement 1). The most interesting feature of the (MmpS4)_3_-(MmpL4)_3_ model is the CCD of MmpL4 which extends along the three-fold symmetry axis 130 Å far into the periplasm. MmpS4 is predicted to be anchored in the cytoplasmic membrane via an N-terminal single transmembrane helix. If we follow the polypeptide sequence of one MmpS4 protomer in the AlphaFold2 model, the transmembrane helix of MmpS4 associates with the 12-transmembrane helix bundle of the first MmpL4 protomer (teal in Figure 1A) by interacting with transmembrane helix 8. A linker devoid of secondary structures bridges the crevice formed between the first (teal) and second (white) MmpL4 protomer, thereby establishing interactions with both MmpL4 protomers. The C-terminal domain of MmpS4 features the topology of a β-sandwich [8, 24], wherein two β-sheets each comprising four β-strands each stack against each other. This architecture is further reinforced by a disulfide bond connecting that connects the sheets. The MmpS4 β-sandwich domain establishes extensive contacts with the CCD of MmpL4, especially the α-helical hairpins donated from the second (white) and the third (grey) MmpL4 protomer. In summary, MmpS4 is predicted to establish extensive contacts with all three MmpL4 protomers, thereby stabilizing the MmpL4 trimer, and in particular the CCD.

**Figure 1.**
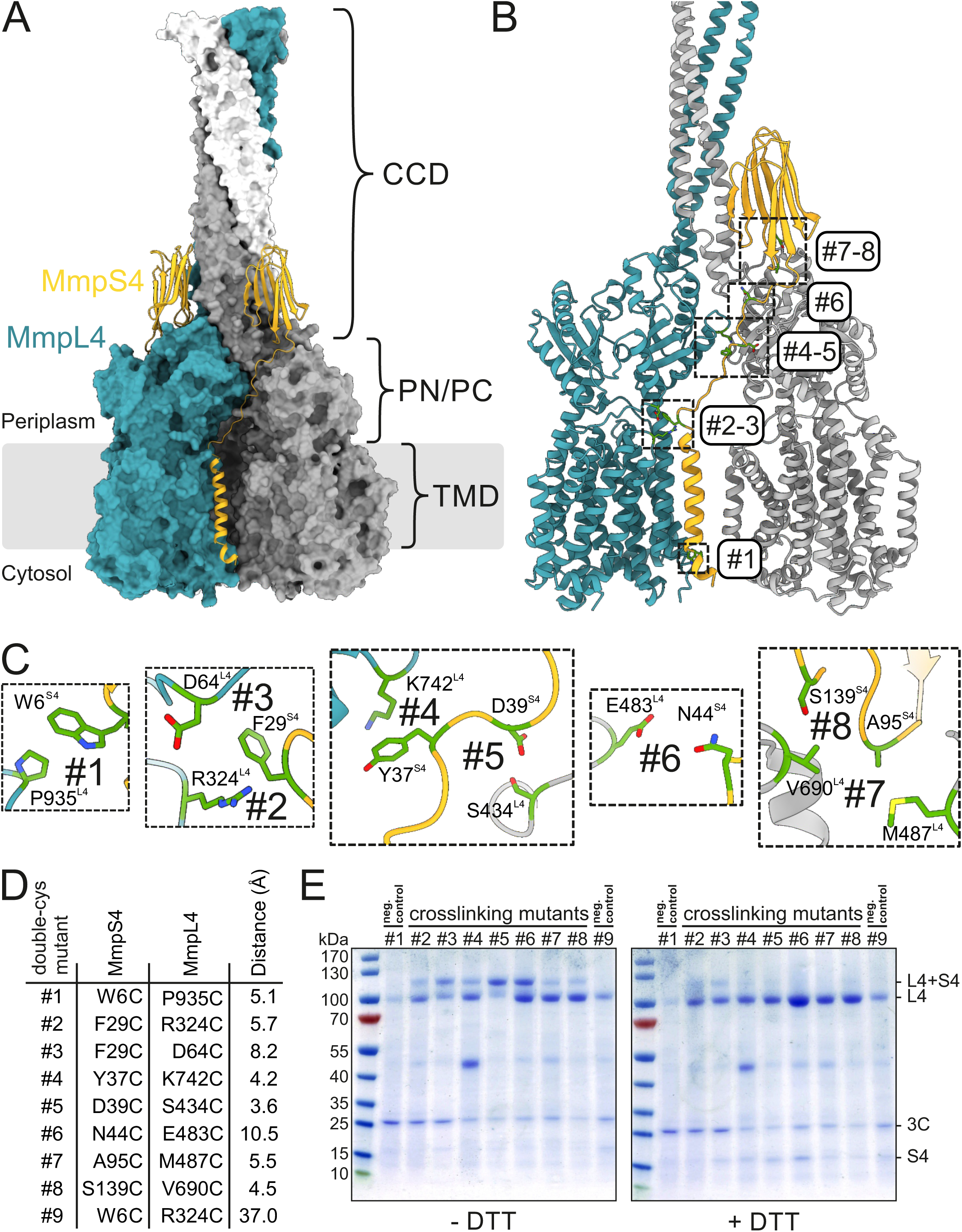
Engineered MmpS4-MmpL4 disulfide cross-links based on AlphaFold2 prediction. (**A**) AlphaFold2 prediction of the hexameric (MmpS4)_3_-(MmpL4)_3_ complex. The three protomers of MmpL4 are shown in surface presentation and colored teal, grey and white. The MmpS4 protomers are shown as yellow cartoon. The boundary of the cytoplasmic membrane is indicated by grey lines. The locations of the transmembrane domains (TMD) and the periplasmic PN/PC domains are indicated. MmpL4 is predicted to form a large coiled-coil domain (CCD) extending into the periplasm. (**B** and **C**) Based on the AlphaFold2 model, eight double-cysteine mutants (#1-#8) were generated, whose thiol groups are in close distance between MmpL4 and MmpS4. The original residues that were substituted by cysteines are shown as sticks. The mutants #2-#8 are expected to form spontaneous disulfide cross-links in the oxidative environment of the periplasm. (**D**) List of characterized double cysteine mutants. Whenever possible, the distances were measured between the γ-atoms of the respective residues of MmpS4 and MmpL4. In case of alanine and proline residues, the atom nearest to the γ-atom of the partner residue was used to measure the distance. (**E**) Non-reducing (left) or reducing (right) SDS-PAGE analysis of purified cysteine cross-linking mutants of MmpS4-MmpL4, stained by coomassie. The negative control cross-linking pair #1 was not expected to form a disulfide bond because it is located in the reducing environment of the cytosol. The negative control cross-linking pair #9 bears cysteines that are too far apart to form a disulfide bond. 3C stands for 3C protease, which was used to elute the complexes from GFP-trap resin.

### Experimental validation of the AlphaFold2-predicted MmpS4-MmpL4 complex

We used targeted disulfide cross-linking to experimentally validate the AlphaFold2-predicted MmpS4-MmpL4 complex. Spontaneous disulfide bonds are expected to form in the oxidative environment of the periplasm of *E. coli* if cysteine pairs are sufficiently close to each other, especially if they are placed in flexible loop regions. The same experimental strategy was previously utilized to study the conformational cycling of the RND transporter AcrB [25], but also more recently to validate the MmpS5-MmpL5 complex structure [16]. To avoid unspecific reactions with free thiol groups, we mutated five out of the seven naturally occurring cysteines of MmpL4 into serines (see methods). We decided to keep two highly conserved cysteine-pairs that form disulfide bonds, the first one at the tip of the CCD of MmpL4 and the second one between the β-sheets of MmpS4.

Based on the AlphaFold2 prediction we introduced eight cysteine pairs (numbered #1-#8) with close distances between the thiol groups into flexible loops along the MmpS4-MmpL4 interface (Figure 1B and C). Cysteine pair #1 located at the cytosol served as a negative control, because it is not expected to form spontaneous disulfide bonds due to the reducing environment. Cysteine pair #9 was included as an additional negative control, because its thiols are 37 Å away from each other and therefore cannot form disulfide bonds (Figure 1D). The respective proteins were produced in *E. coli*. Prior to the detergent solubilization step, n-ethylmaleimide was added to react with free reactive thiols and prevent undesired disulfide cross-links between transporter complexes during detergent extraction and subsequent purification steps. Subsequently, the transporter complexes were purified via GFP attached C-terminally to MmpL4 and analyzed by non-reducing and reducing SDS-PAGE (Figure 1E). Cysteine pairs #2-8 established variable degrees of MmpS4-MmpL4 cross-linking. As expected, the cytoplasmic pair #1 and the negative control pair #9 did not cross-link, thereby validating the experimental approach. The degree of cross-linked protein for cysteine pair #5 (D39C^MmpS4^-S434C^MmpL4^), which according to the AlphaFold2 prediction has the closest distance of 3.6 Å between the thiol moieties, was the highest. In summary, the engineered cysteine cross-links provided experimental evidence for the interaction between and trimerization of MmpS4 and MmpL4 as predicted by the AlphaFold2 model.

### Cross-linked MmpS4-MmpL4 complexes are functionally impaired

Next, we assessed the functional impact of the cysteine pairs #3, #5 and #8 (Figure 2). To this end, we used a previously established toxicity assay, which builds on the observation that cells lacking both mycobactin exporters MmpL4 and MmpL5 are highly susceptible to externally added mycobactin accumulating within the cell [9]. Genes encoding the respective MmpS4-MmpL4 variants were expressed on a plasmid in an *M. tuberculosis* strain lacking both MmpL4 and MmpL5 (*MtbΔmmpL4/mmpL5*). Western blots detecting the C-terminal FLAG_3_-tag revealed that cys-depleted MmpS4-MmpL4 as well as the engineered MmpS4 and MmpL4 cysteine variants were produced at similar levels (Figure 2 — figure supplement 1). Under non-reducing conditions, MmpL4-MmpS4 complexes were detected in denaturing gels for the three cysteine pairs in contrast to the single cysteine mutants, indicating disulfide bond formation upon expression in *M. tuberculosis* (Figure 2 — figure supplement 1). This result is consistent with disulfide crosslinking observed in *E. coli* (Figure 1). For the toxicity assays, cys-depleted MmpS4-MmpL4 was included as a reference, because all cysteine variants were introduced into this background. Of note, cys-depleted MmpS4-MmpL4 exhibited a rather low activity compared to previous complementation experiments with wild-type MmpS4-MmpL4 [15], indicating that the substituted cysteines in MmpL4 contribute to the activity of the MmpL4-MmpS4 complex. The single cysteine variants of the three investigated cysteine pairs were similarly active as the cys-depleted MmpS4-MmpL4 control (Figure 2). In contrast, constructs containing the cysteine pairs #3 and #8, but not cysteine pair #5, were functionally impaired. Interestingly, the strain expressing the cysteine pairs #3 was even more susceptible to mycobactin than *MtbΔmmpL4/mmpL5*. In this regard, it is noteworthy that cysteine cross-link #3 tethers MmpS4 to MmpL4 at the interface of the transmembrane and the periplasmic domainss, while cysteine cross-link #8 blocks the relative movement of the β-sandwich domain of MmpS4 against the periplasmic domain of MmpL4. The observed activity loss upon cross-link formation suggests that these structural elements move relative to each other during the transport cycle. On the other hand, cysteine cross-link #3 is not affected at the functional level, likely owing to the fact that two rather flexible loops were tied together, which allows for conformational flexibility.

**Figure 2.**
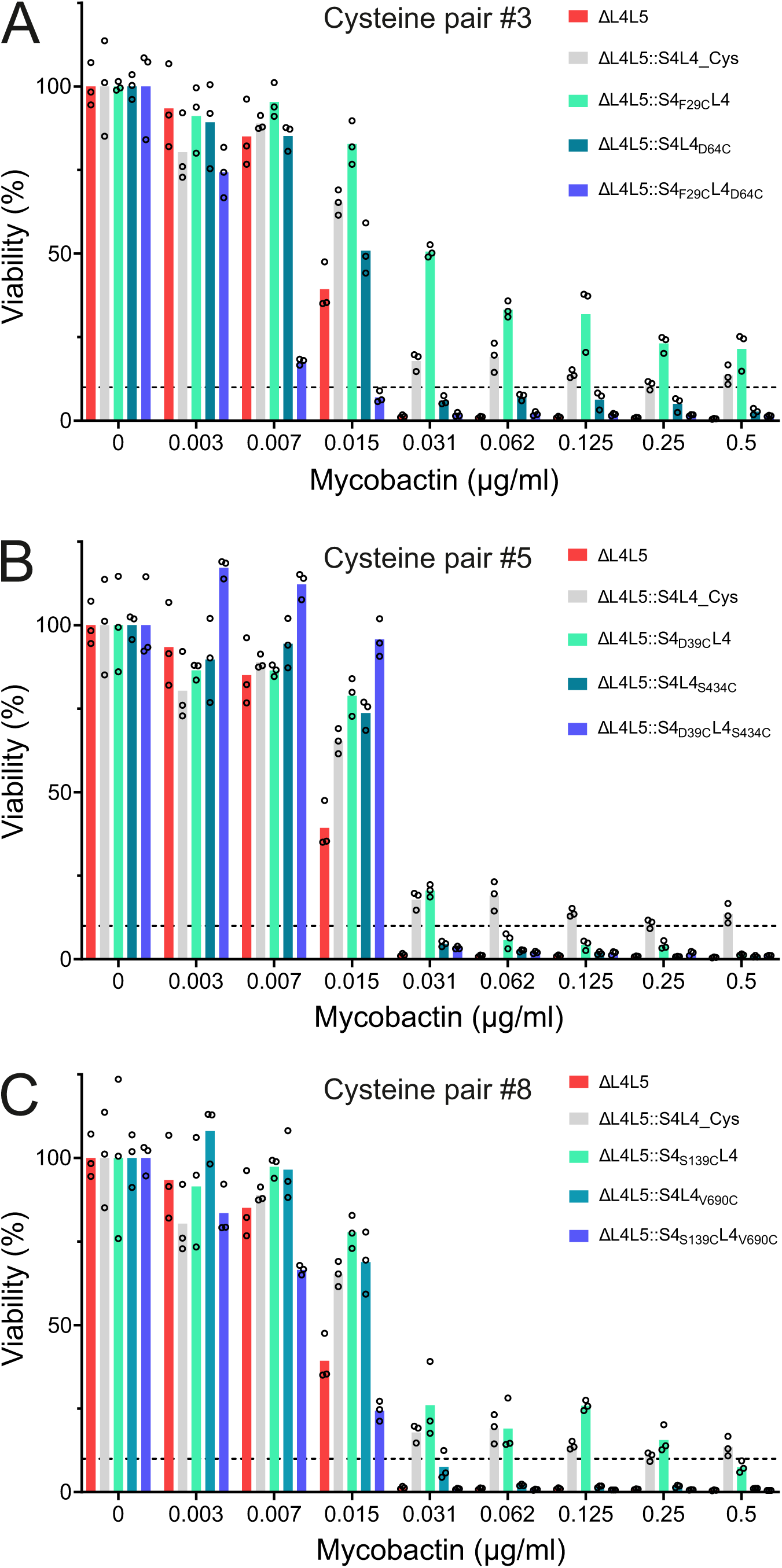
Functional analysis of MmpS4 and MmpL4 cysteine variants. The viability of *M. tuberculosis* strains at increasing mycobactin concentrations was determined using the microplate Alamar Blue assay (see methods). (**A-C**) Functional analysis of cysteine pair #3 (MmpS4_F29C_ - MmpL4_D64C_) (**A**), cysteine pair #5 (MmpS4_D39C_ - MmpL4_S434C_) (**B**) and cysteine pair #8 (MmpS4_S139C_ - MmpL4_V690C_) (**C**). Data were normalized to the viability in the absence of mycobactin (100%). ΔL4L5, *M. tuberculosis* strain lacking *mmpL4* and *mmpL5*; :S4L4_Cys, complementation with cysteine-depleted *mmpS4-mmpL4*; ::S4_F29C_L4 complementation with cysteine-depleted *mmpS4-mmpL4* with the F29C mutation introduced into MmpS4 (and for the other cysteine variants accordingly). Data points correspond to technical triplicates.

### Cryo-EM structure of the (MmpS4)_3_-(MmpL4)_3_ complex

The cysteine cross-linking pair #5, which exhibited the strongest cross-linking efficiency and whose activity was not affected, was purified (Figure 3 — figure supplement 1) and analyzed by single particle cryo-EM, resulting in a high-resolution reconstruction of the hexameric (MmpS4)_3_-(MmpL4)_3_ complex (Figure 3, Figure 3 — figure supplement 2, Figure 3 — figure supplement 3, Table 1). We obtained a 3.1 Å map without applying symmetry during data processing (Figure 3 — figure supplement 2). An additional density for the *E. coli* ACP was present at the cytosolic interface of the transmembrane domain, similar to what has previously been described in the structure of the MmpL4 monomer.

**Figure 3.**
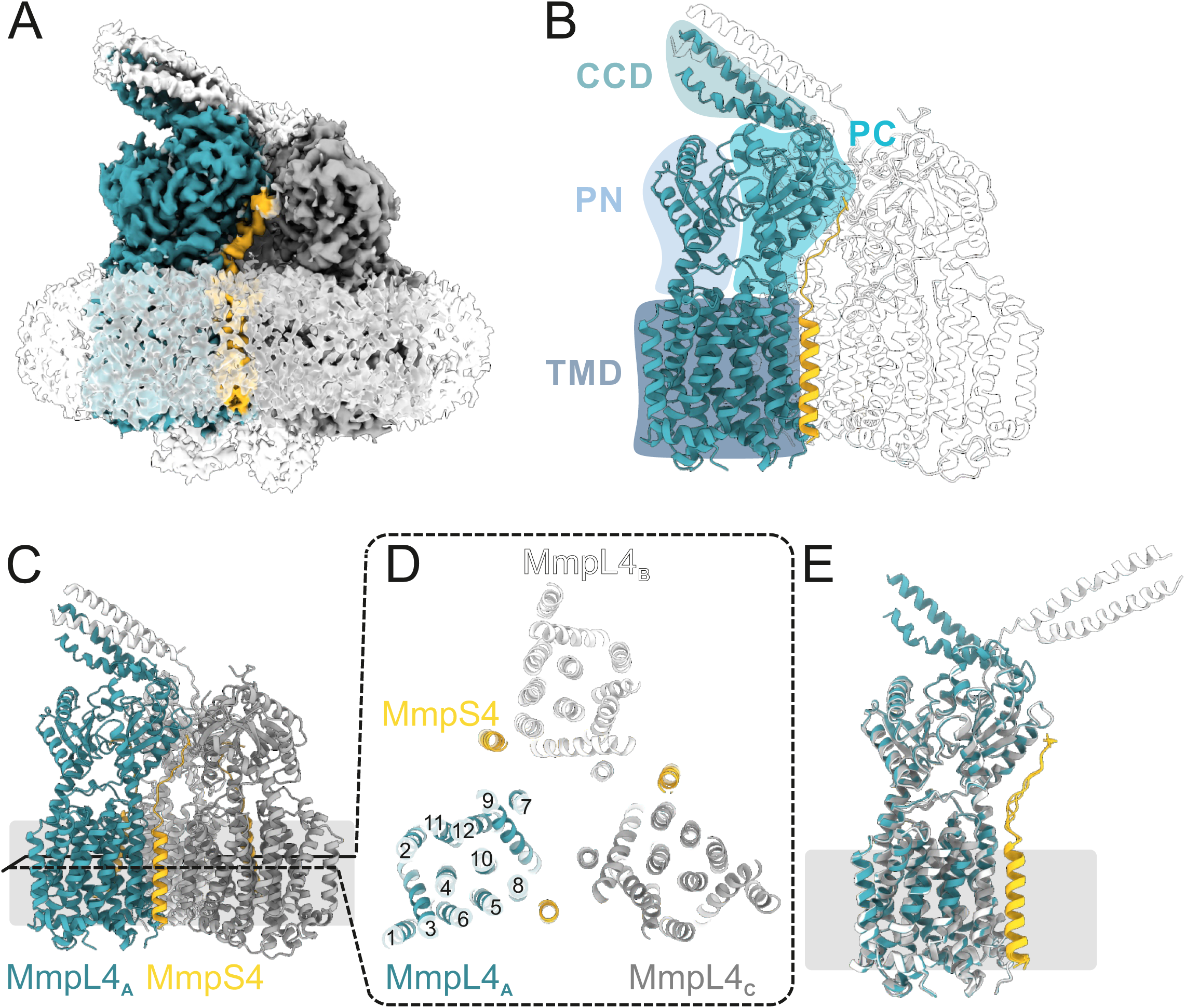
Structure of the hexameric (MmpS4)_3_-(MmpL4)_3_ complex. (**A**) Left panel; Cryo-EM map of the hexameric (MmpS4)_3_-(MmpL4)_3_ complex processed using C1 symmetry viewed along the membrane plane and contoured at 4σ. The detergent micelle and weak density for *E. coli* ACP are shown as white, transparent density. MmpL4 protomers are colored in teal, grey and white, MmpS4 in yellow. Density for a tilted CCD formed out of two α-helical hairpins donated from MmpL4 protomers A (teal) and B (white) is visible. (**B**) Schematic labeling of MmpL4 domains. (**C**) Structural model of the asymmetric (MmpS4)_3_-(MmpL4)_3_ complex shown as cartoon shown along the membrane plane (left). (**D**) Transmembrane helices of MmpL4 and MmpS4 viewed from the periplasm. The transmembrane helices of MmpL4 are numbered. (**E**) Superimposed MmpS4-MmpL4 monomers of asymmetrically processed (MmpS4) - (MmpL4) complex.

**Table 1:**
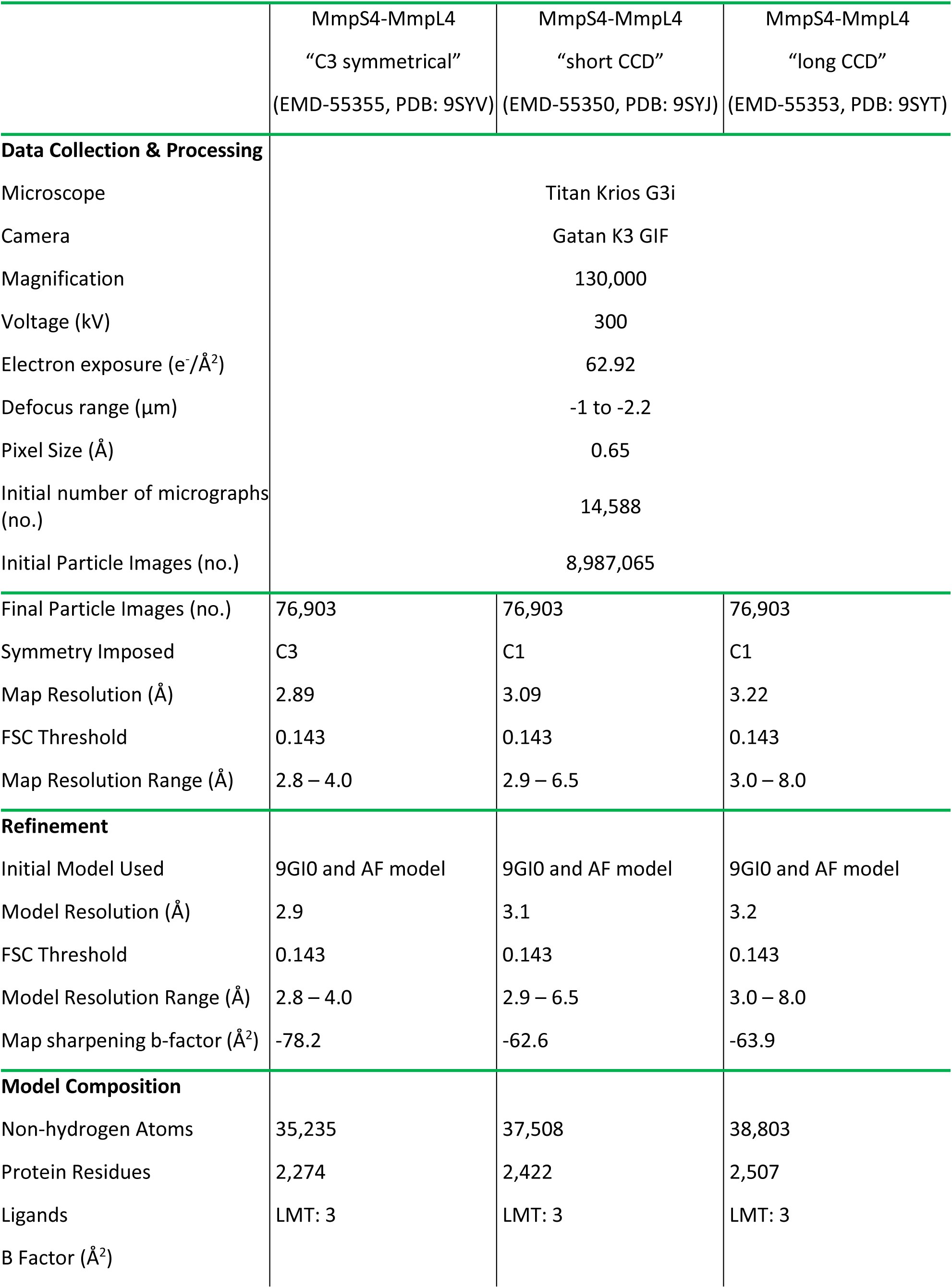
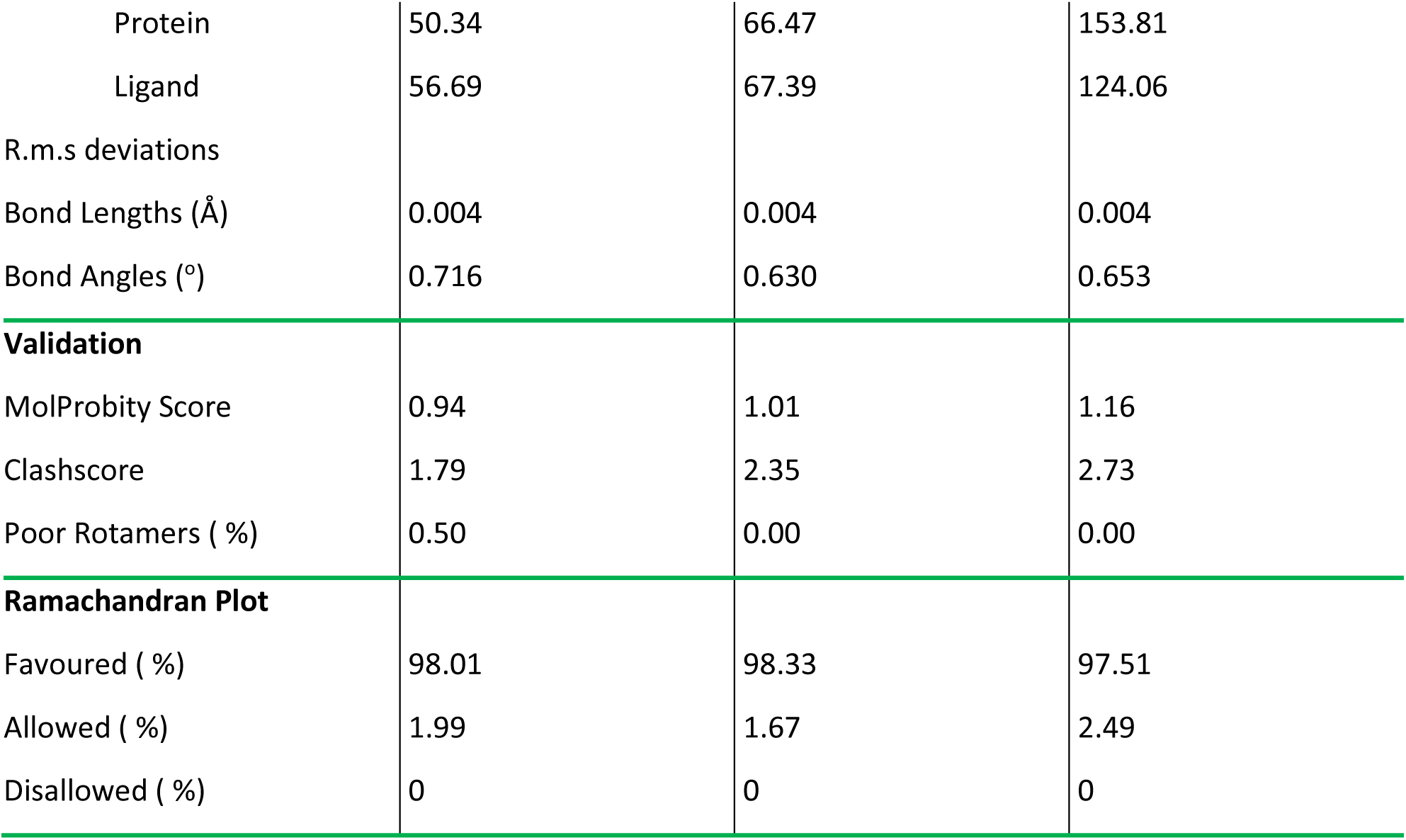
Cryo-electron microscopy data collection statistics.

|  | MmpS4-MmpL4<br>“C3 symmetrical”<br>(EMD-55355, PDB: 9SYV) | MmpS4-MmpL4<br>“short CCD”<br>(EMD-55350, PDB: 9SYJ) | MmpS4-MmpL4<br>“long CCD”<br>(EMD-55353, PDB: 9SYT) |
|---|---|---|---|
| **Data Collection & Processing** |  |  |  |
| Microscope | Titan Krios G3i |  |  |
| Camera | Gatan K3 GIF |  |  |
| Magnification | 130,000 |  |  |
| Voltage (kV) | 300 |  |  |
| Electron exposure (e<sup>-</sup>/Å<sup>2</sup>) | 62.92 |  |  |
| Defocus range (µm) | -1 to -2.2 |  |  |
| Pixel Size (Å) | 0.65 |  |  |
| Initial number of micrographs (no.) | 14,588 |  |  |
| Initial Particle Images (no.) | 8,987,065 |  |  |
| Final Particle Images (no.) | 76,903 | 76,903 | 76,903 |
| Symmetry Imposed | C3 | C1 | C1 |
| Map Resolution (Å) | 2.89 | 3.09 | 3.22 |
| FSC Threshold | 0.143 | 0.143 | 0.143 |
| Map Resolution Range (Å) | 2.8 – 4.0 | 2.9 – 6.5 | 3.0 – 8.0 |
| **Refinement** |  |  |  |
| Initial Model Used | 9GI0 and AF model | 9GI0 and AF model | 9GI0 and AF model |
| Model Resolution (Å) | 2.9 | 3.1 | 3.2 |
| FSC Threshold | 0.143 | 0.143 | 0.143 |
| Model Resolution Range (Å) | 2.8 – 4.0 | 2.9 – 6.5 | 3.0 – 8.0 |
| Map sharpening b-factor (Å<sup>2</sup>) | -78.2 | -62.6 | -63.9 |
| **Model Composition** |  |  |  |
| Non-hydrogen Atoms | 35,235 | 37,508 | 38,803 |
| Protein Residues | 2,274 | 2,422 | 2,507 |
| Ligands | LMT: 3 | LMT: 3 | LMT: 3 |
| B Factor (Å<sup>2</sup>) |  |  |  |
| Protein | 50.34 | 66.47 | 153.81 |
| Ligand | 56.69 | 67.39 | 124.06 |
| R.m.s deviations |  |  |  |
| Bond Lengths (Å) | 0.004 | 0.004 | 0.004 |
| Bond Angles (°) | 0.716 | 0.630 | 0.653 |
| **Validation** |  |  |  |
| MolProbity Score | 0.94 | 1.01 | 1.16 |
| Clashscore | 1.79 | 2.35 | 2.73 |
| Poor Rotamers ( %) | 0.50 | 0.00 | 0.00 |
| **Ramachandran Plot** |  |  |  |
| Favoured ( %) | 98.01 | 98.33 | 97.51 |
| Allowed ( %) | 1.99 | 1.67 | 2.49 |
| Disallowed ( %) | 0 | 0 | 0 |

Because this density was only weakly defined (Figure 3A, Figure 3 — figure supplement 2), we did not include it in our model. The C1 cryo-EM map contained well-defined density for the transmembrane helices and part of the flexible linkers of MmpS4, but not for its β-sandwich domains (Figure 3A, Figure 3 — figure supplement 3). Well-resolved density is present for the transmembrane domains and the periplasmic PN and PC domains of MmpL4, whereas the CCD is less well defined (Figure 3A and B). Strikingly, the CCD is tilted by around 60° away from the threefold symmetry axis rather than extending along it as predicted by AlphaFold2. Only two out of three α-helical hairpins of the three MmpL4 protomers are resolved (Figure 3C), which may explain its tilted conformation. Using a local refinement approach with a mask containing the CCD and the uppermost regions of MmpL4, the cryo-EM density of the CCD and the loops connecting to the PC domain of MmpL4 was further improved (Figure 3 — figure supplement 2, Figure 3 — figure supplement 3), enabling us to build a model of the lower part of the CCD, including the linkers connecting to the PC of MmpL4. 3D flexible refinement performed with CryoSPARC further supports the notion that the tilted CCD is rather flexible (Figure 3 — video 1). Model building revealed that the tilted CCD is composed of the α-helical hairpins of the polypeptide chains A and B. To assess whether there are asymmetries in other parts of the structure, we superimposed the individual protomers of the (MmpS4)_3_-(MmpL4)_3_ complex analyzed using C1 symmetry (Figure 3E). Apart from the two resolved α-helical hairpins, the MmpL4 core domains and the resolved parts of MmpS4 differ by a RMSD of less than 0.6 Å and are therefore structurally identical considering the map resolution of around 3 Å. To increase the resolution of the core parts of the (MmpS4)_3_-(MmpL4)_3_ complex, we applied C3 symmetry resulting in a map with a resolution of 2.9 Å, but lacking density for the CCD (Figure 3 — figure supplement 2). Using this data, we built a model comprising residues E16-M487 and V690-R939 of MmpL4 and residues T5-T40 of MmpS4.

### Identification of a deep cavity accommodating a detergent molecule

A systematic inspection of non-proteinaceous densities revealed a strong detergent-like density in a cavity formed between the PN and PC domains and the upper part of the TMD (Figure 4A and B, Figure 4 — figure supplement 1). We modeled the density as a n-dodecyl β-D-maltoside (β-DDM) molecule, the detergent used to extract and purify the transporter complex. The two sugar moieties are sandwiched between the PN and PC and the twelve carbon atoms of the aliphatic chain are accommodated in a hydrophobic patch in the transmembrane domain (Figure 4B). We did not observe density for β-DDM at this location in the monomeric MmpL4 structure, although β-DDM was also used for protein extraction and purification. Using the 3 V tool [26], we analyzed the cavity that accommodates β-DDM in the (MmpS4)_3_-(MmpL4)_3_ complex and found that it spans from the TMD into the interface between PN and PC (Figure 4D). The cavity does not open up to the periplasm as observed for the open conformation of MmpL3 (Figure 4E). Instead, an entrance is found between transmembrane helices 1 and 2, at the same position at which Fountain et al. observed a prominent non-proteinaceous density in the MmpS5-MmpL5 complex structure [21]. In striking contrast, monomeric MmpL4 only contains a very small, occluded cavity at the same location (Figure 4C). We also noted extra-densities wedged between the transmembrane helix bundles at the inner leaflet of the membrane (Figure 4 — figure supplement 1), in analogy to the lipids observed in the MmpS5-MmpL5 structures that were suggested to stabilize the MmpL5 trimer [16, 21]. Because they were not sufficiently defined to unambiguously differentiate between lipid or detergent molecules we refrained from interpreting them. It should be noted that we purified the MmpS4-MmpL4 complex from *E. coli* as expression host, whereas the MmpLS5-MmpL5 complexes were purified from *M. smegmatis*.

**Figure 4.**
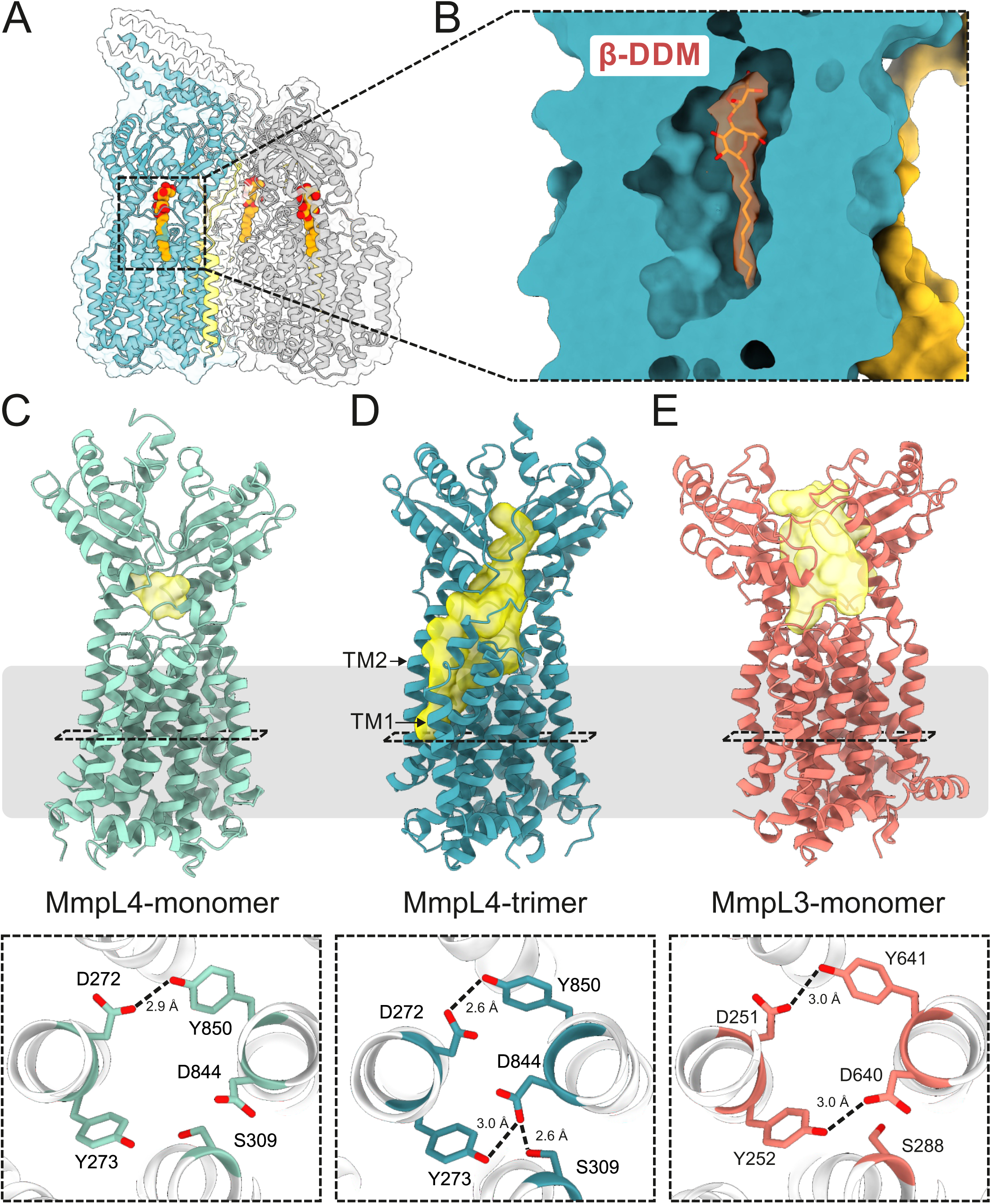
Detergent binding in a deep cavity extending into the periplasmic domain of MmpL4. (**A**) Structural model of hexameric (MmpS4)_3_-(MmpL4)_3_ complex processed using C1 symmetry. Three detergent molecules (β-DDM) bound to each of the MmpL4 protomers are depicted as red/orange spheres. (**B**) Enlarged view of the β-DDM binding site of chain A. Non-proteinaceaous density (contoured at 6.8σ) interpreted as β-DDM is depicted in orange. (**C** – **E**) Cavity analysis of monomeric MmpL4 (PDB ID: 9GI0) (**C**), trimeric MmpL4 (this work, PDB ID: 9SYJ) (**D**) and monomeric MmpL3 (PDB ID: 7NVH) (**E**) using the 3V tool. Cavities are shown in yellow and the membrane boundary is indicated. The bottom row shows the conserved DY-pairs and S309(MmpL4)/S288(MmpL3) viewed from the periplasm. Hydrogen bonds are indicated by dashed lines.

### Comparison of monomeric and trimeric MmpL4 structures

To systematically analyze the conformational differences between monomeric and multimeric MmpL4, we used the Needleman–Wunsch algorithm implemented in ChimeraX to superimpose the 3.0 Å MmpL4 monomer structure (PDB ID: 9GI0) onto the first MmpL4 protomer (chain A) of the asymmetric (MmpS4)_3_-(MmpL4)_3_ complex structure determined at 3.1 Å (Figure 5A and B). The RMSD was 2.14 Å over 722 residues, demonstrating that MmpL4 exhibits substantial structural differences depending on whether it is a monomeric protein or embedded in the hexameric complex. When plotting the C_α_-atom distance of monomeric versus trimeric MmpL4 (Figure 5C), we found that conformational differences affect all parts of the transporter. Most prominently, we observed a rotation of the PN domain (Figure 5C), which is responsible for creating the deep cavity accommodating β-DDM in the complex structure (Figure 4A and B). Intriguingly, we also observed a rearrangement of the conserved DY-pairs, four key residues residing at the center of the transmembrane domain that are suggested to couple active transport to the proton-gradient [15] (Figure 4C and D, Figure 4 — figure supplement 1). While Y273 and D844 are around 5 Å apart in monomeric MmpL4 (Figure 4C), they engage in a hydrogen bond in the context of the hexameric complex (Figure 4D) in a similar configuration as observed for MmpL3 (PDB ID: 7NVH) (Figure 4E). These rearrangements may be coupled to protonation events at D844 in a similar manner as characterized for the proton translocating aspartates in the drug efflux pump AcrB [27]. Concomitant with the subtle rearrangements in the proton-coupling DY-pairs, the twelve transmembrane helices of MmpL4 change their position (RMSD of 1.78 Å over 369 residues) to a similar extent as the conformational changes observed for the TMD of asymmetric AcrB [28]. In MmpL4, the conformational changes mostly affect the upper parts of the transmembrane helices, in particular transmembrane helix 2-4 and 11-12 (Figure 5B and C). These conformational changes result in a larger distance between transmembrane helices 1 and 2, creating the channel entry that leads to the periplasmic cavity (Figure 4D). It remains to be shown in future work whether the structural differences observed between monomeric and multimeric MmpL4 represent alternate conformational states that are relevant for efflux function.

**Figure 5.**
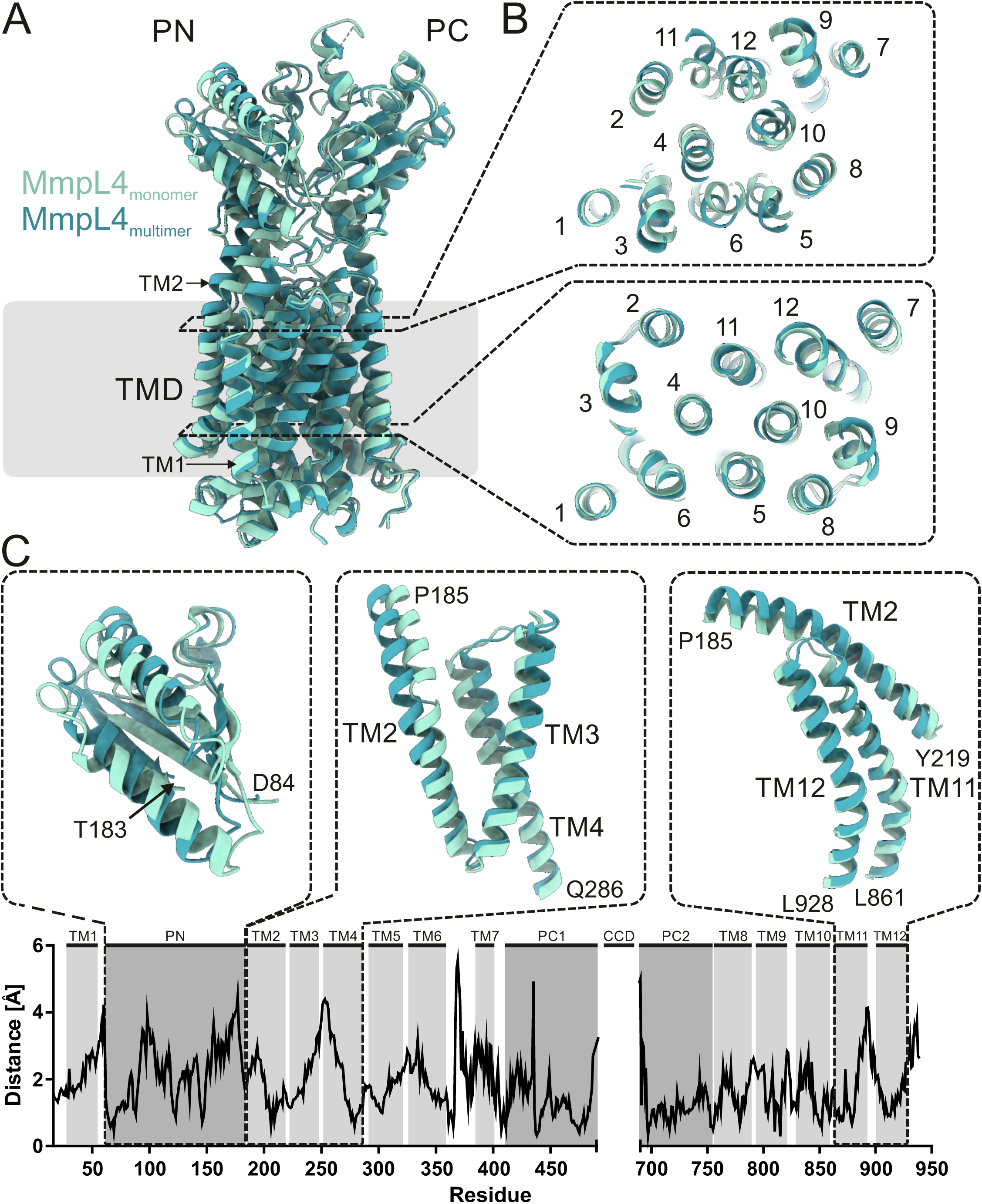
Systematic structural comparison of monomeric versus trimeric MmpL4. (**A** and **B**) Previously determined monomeric MmpL4 colored in light blue (PDB ID: 9GI0) was superimposed onto trimeric MmpL4 colored in dark blue (chain A of structure obtained using C1 symmetry, PDB ID: 9SYJ). (**B**) Two slices along the membrane plane are shown from the periplasm onto the TMDs at the two heights indicated by the dashed rectangles in (**A**). (**C**) C_α_-atom distance plot between monomeric and trimeric MmpL4. The respective TM helices and domains are indicated above. For three structural elements showing marked differences, the superimposed structures are shown as cartoon above the plot. No structural information was available for the CCD and the C-terminus. The overall RMSD is 2.14 Å over 722 residues.

### Structural features of MmpL CCDs analyzed by AlphaFold

The presence of defined density for the tilted CCD consisting of α-helical hairpins of only two MmpL4 protomers suggests that it represents a thermodynamically stable intermediate. The density for the CCD was less well defined than other parts of MmpL4, suggesting that the CCD is rather flexible (Figure 3 — figure supplement 3). To gain further insights into this unique structural element, we used AlphaFold3 to predict the CCDs of all cluster I MmpL proteins of *M. tuberculosis* and performed a sequence analysis of this region (Figure 6 — figure supplement 1). In terms of phylogeny, Cluster I MmpL proteins can be further subdivided into two groups, the first one encompassing MmpL1, MmpL2, MmpL4, MmpL5, MmpL6 and MmpL9 (which apart from MmpL9 are encoded in an operon with a respective *mmpS* gene) and the second one encompassing MmpL8, MmpL10 and MmpL12 (Figure 6 — figure supplement 1). MmpL7 cannot be clearly assigned to either of the groups. The predicted CCDs differ quite strikingly between the two phylogenetic groups, with the second group (MmpL8, MmpL10 and MmpL12) being substantially longer and containing kinks and discontinuities in the α-helices. It should be noted that the AlphaFold3 pLDDT values were less confident for MmpL7, MmpL8, MmpL10 and MmpL12 as compared to the other MmpLs, and future experimental structures will be required to validate their correctness.

The most notable feature of the CCDs is the enrichment of methionine side chains pointing towards the enclosed channel for many, but not all MmpL proteins (Figure 6A, Figure 6 — figure supplement 1). The channel wall of MmpL4’s CCD, for example, is lined with 45 methionines (15 per α-hairpin). By contrast, MmpL6 has one methionine per α-hairpin, suggesting that the methionines are not *per se* needed for the assembly of the CCD, but instead likely play a functional role during transport.

**Figure 6.**
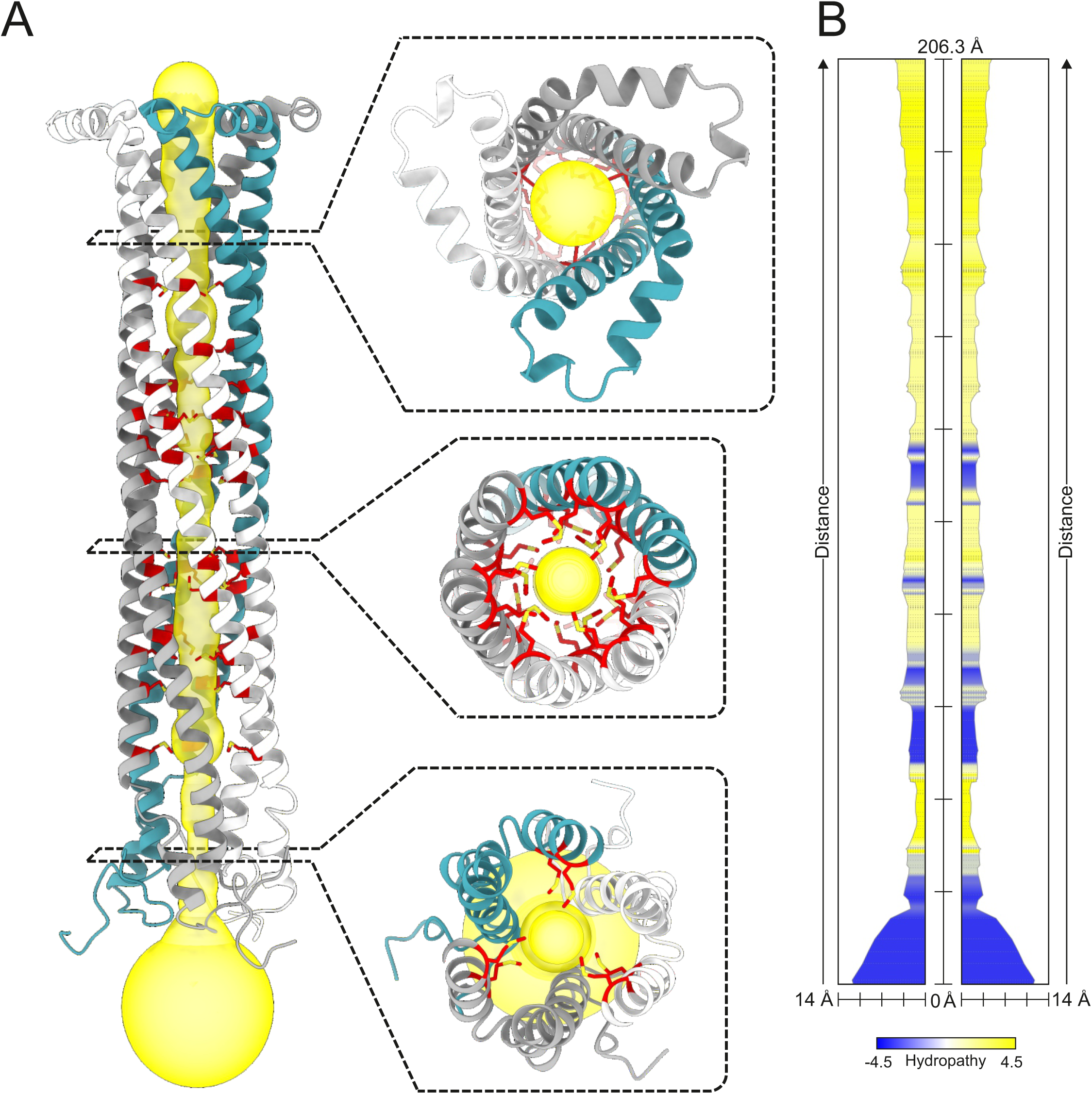
Channel prediction of the coiled-coil domain of MmpL4. (**A**) The CCD of trimeric MmpL4 as predicted by AlphaFold3 and shown as cartoon. A continuous channel leading through the CCD interior was calculated by the MOLEonline tool and shown as yellow surface. Methionine side chains lining the channel surface are shown as red sticks. Three top views along the pore are shown as insets. (**B**) Radius and biophysical properties of the channel shown in (**A**) calculated by the MOLEonline tool.

The CCD channel radius of all MmpL proteins appears to be very narrow. The MOLEonline tool [29] predicts several constrictions and a minimal channel radius of 1.1 Å for the CCD of MmpL4 (Figure 6B). It should be kept in mind that these analyses were performed with AlphaFold predictions and not with experimental structures. In its closed conformation, the channel appears to be too narrow to let bulky substrates such as desferrated mycobactin or bedaquiline pass through. However, flexibility of the methionine side chains coating the channel wall as well as the overall conformational flexibility of the CCD might nevertheless provide sufficient dilation to let transport substrates pass. Further experimental studies will therefore be required to deepen our mechanistic understanding of the CCD, in particular with regards to its role in substrate transport.

## DISCUSSION

Structure predictions using AlphaFold have become a standard step in every structural biology project, due to its ease of use and the high accuracy of its predictions [23]. Predictions of multi-protein complexes by AlphaFold2 multimer (and more recently AlphaFold3) are especially attractive, because they provide potential cues about the interaction interfaces as well as possible multimeric assemblies [30, 31]. In the context of MmpL and MmpS proteins, AlphaFold has been extensively used by us and others to firstly identify the CCD in MmpL4 and MmpL5 and to secondly predict its involvement in trimerization of the proteins [15, 16, 18, 21]. In our previous study on MmpL4, we cloned a transporter construct devoid of the predicted CCD to enable the determination of a highly resolved cryo-EM structure of the MmpL4 monomer [15]. A similar strategy was also chosen by Fountain et al. to obtain the multimeric MmpS5-MmpL5 structure [21]. In our study, we used the hexameric structure of MmpS4-MmpL4 predicted by AlphaFold2 as the basis to introduce engineered cysteine cross-links. This approach enabled us to biochemically validate the predicted model and facilitated its structural determination. The procedure described in this work may therefore serve as a blueprint on how AlphaFold predictions can aid the rational stabilization of difficult-to-handle protein complexes. Other groups have determined cryo-EM complex structures of the MmpS4-MmpL4 homologues MmpS5-MmpL5 [16, 21] and MSEMG_1381-1382 [20]. These structures were determined without cross-linking, either by purifying the complexes from *M. smegmatis* [16, 21] (instead of *E. coli* as in this study) or by using grid preparation methods that avoid purification of the membrane protein complexes after solubilization [20]. Independently from the sample preparation approach, hexameric complexes composed of three MmpS and three MmpL proteins were observed, demonstrating that cross-linking via engineered cysteines did not result in structural artefacts. In addition, the MmpS4_3_-MmpL4_3_ complex we obtained is structurally very similar to the MmpL5 structures obtained by the other two groups (RMSD values smaller than 1.4 Å for the core domains of MmpL4/MmpL5) (Figure 3 — figure supplement 4).

AlphaFold predictions and structures of the multimeric assembly of MmpS-MmpL proteins presented in this study and by others [16, 20, 21] provide important insights into the architecture of these unique efflux machineries. The structures established the role of the MmpS protein in stabilizing the MmpL trimer by interactions of every MmpS protomer with all three MmpL protomers. We did not observe density for the β-sandwich domains of MmpS4, nor were the respective domains of MmpS5 observed in the symmetric hexameric MmpS5-MmpL5 complex [16]. Our cross-linking data, however, strongly suggests that the β-sandwich domains of MmpS4 interact with the CCD in the configuration predicted by AlphaFold2 (see cross-links of cysteine pairs #7 and #8 in Figure 1A) and this was also confirmed by the cryo-EM structure of MSEMG_1381-1382 [20]. We therefore hypothesize that the role of the β-sandwich domains of MmpS proteins is to further stabilize the fully assembled CCD in its upright position and orchestrate its orientation in the periplasm (Figure 7).

**Figure 7.**
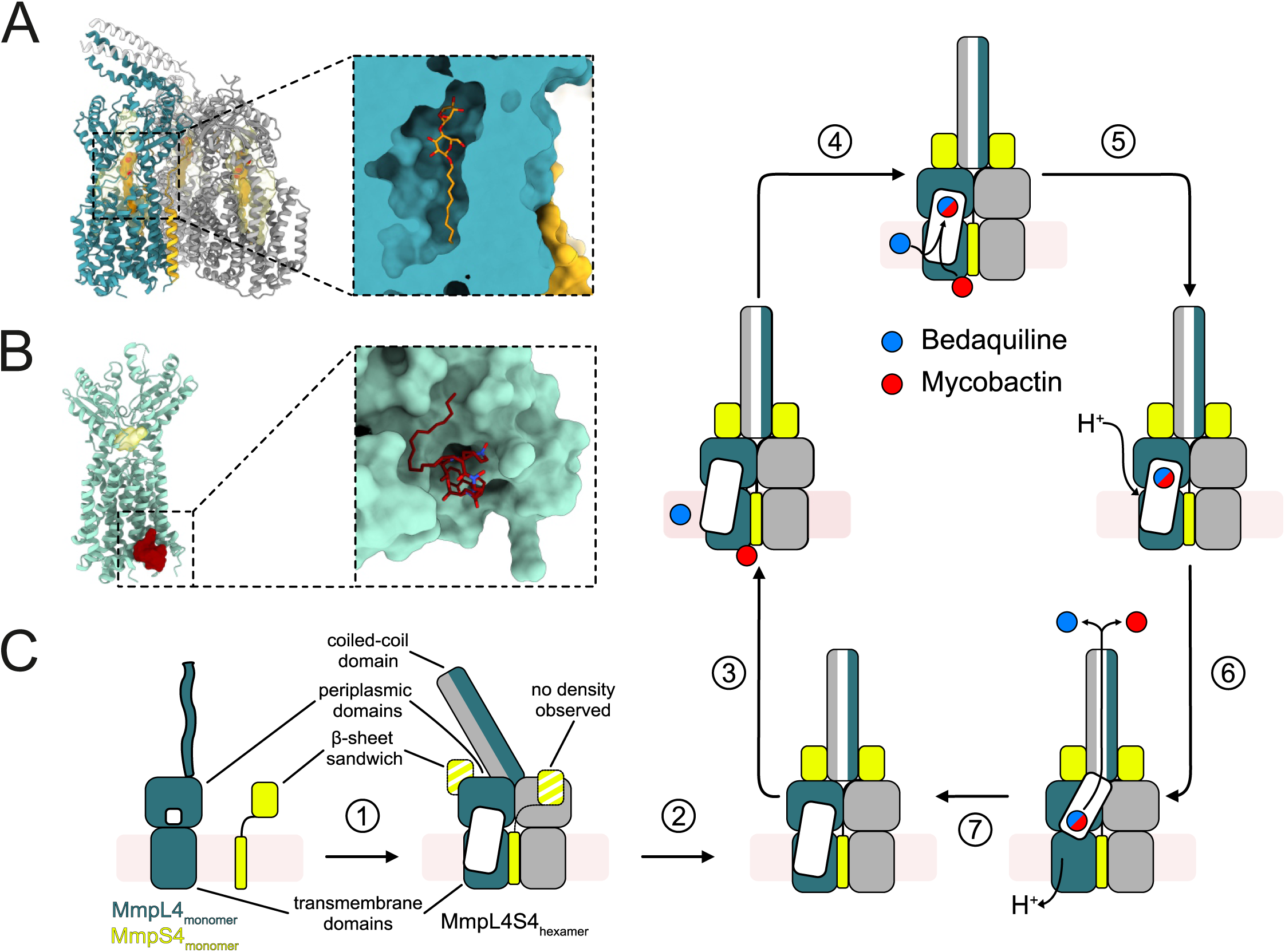
Mycobactin and bedaquiline export mechanism in *M. tuberculosis*. (**A** and **B**) Hexameric MmpS4-MmpL4 with bound β-DDM (PDB: 9SYJ, this work) (**A**) and monomeric MmpL4 with bound mycobactin (PDB ID: 9GI0, previous study) (**B**), representing the two experimentally determined cryo-EM structures of MmpL4. (**C**) Proposed mechanism of transport. ①: Experimentally validated step wherein MmpS4 (yellow) and MmpL4 (teal, grey and white) multimerize to form a hexamer. Thereby, a periplasmic cavity (white) accessible from the outer leaflet of the inner membrane is created. The partially assembled CCD is tilted. MmpS4’s β-sheet sandwich is flexible and does not engage in binding. Step ②: Hypothetical step supported by AlphaFold predictions and structures of other homologues. The tilted CCD fully assembles and becomes upright, thereby extending along the three-fold symmetry axis of the transporter complex. MmpS4’s β-sheet sandwich domains engage with the CCD. ③ and ④: Bedaquiline (BDQ) or mycobactin (MBT) are proposed to enter the periplasmic binding cavity either directly from the outer membrane leaflet (BDQ) or from its initial cytoplasmic binding site (MBT). ⑤ and ⑥: Hypothetical steps proposing that protonation events at the transmembrane domains invoke a conformational switch of the periplasmic substrate binding channel to a assume a conformation similar to the one observed for MmpL3. Thereby, bound BDQ and MBT are transported towards the CCD. ⑦: Hypothetical step proposing that deprotonation of the DY-pair resets the transport system to accommodate the next substrate.

The CCDs of cluster I MmpL proteins are particularly interesting from a structural and functional point of view. In their fully assembled, upright and trimeric conformation, the CCDs protrude into the periplasm [15] [16, 20, 21] (Figure 7). A length estimate based on the cryo-ET analysis of the mycobacterial cell membranes suggested that the upright CCD does not bridge the entire periplasmic space of *M. tuberculosis* [21], and therefore it is likely that further protein partners, likely PE/PPE proteins located in the outer membrane [32], are needed to complete the bridge. In the hexameric (MmpS4)_3_-(MmpL4)_3_ complex structure presented in this work, the CCD adopts a partially assembled, tilted conformation, consisting of only two α-helical hairpins. High symmetry among the MmpL4 core protomers (Figure 3E) speak against the tilted CCD conformation to be a cross-linking artefact, though it can not be excluded. Currently, we do not have experimental evidence showing that the tilted CCD has any physiological role, and we therefore suggest that in order to achieve transport-competence, the CCD has to assume its upright, fully assembled conformation (Figure 7). Next to the β-sandwich domains of MmpS4, it is likely that additional periplasmic proteins are required to position the CCD into its transport-competent conformation, as for example Rv0455c which was shown to be the only single protein essential for mycobactin export by interacting with both MmpS4-MmpL4 and MmpS5-MmpL5 export systems [33].

A general finding from our and other studies is the overall flexibility of the CCD. In our structure, the density for the tilted CCD is less defined than other parts of the transporter (Figure 3 — figure supplement 3). Similarly, the CCD of MmpL5 was poorly defined in the MmpS5-MmpL5 complex structure determined with full-length MmpL5 [16], while in the study of Fountain et al., the CCD of MmpL5 was genetically truncated [21]. In contrast, the MSEMG_1381-1382 complex structure revealed a fully assembled, upright CCD and a clear density for the β-sandwich domains of the MmpS protein intimately interacting with the CCDs, as predicted by AlphaFold2 [20], which may be explained by a more robust CCD of MSEMG_1382 compared to MmpL4 and MmpL5 or additional stabilization by the β-sandwich domains of MSMEG_1381.

An important open research questions concerns the substrate pathway through MmpL4 and how it is coupled to the proton motive force. A key experimental finding of this study is the formation of a periplasmic substrate binding cavity extending from a groove formed by TM1 and TM2 in MmpL4 (with β-DDM bound) as a consequence of hexamer formation together with MmpS4 (Step 1 in Figure 7). A similar cavity was observed in multimeric MmpL5 [21]. Based on mutagenesis experiments performed by Fountain et al. [21], there is solid experimental evidence that this cavity plays a role for bedaquiline transport and, by extension, likely also mycobactin transport. Interestingly, in AcrAB-TolC, multiple cavities and transport routes exist [34–36]. Fusidic acid, β-lactams, and β-DDM have been proposed to use a channel extending from a groove formed by TM1 and TM2 to reach their binding pocket [36], hence at an analogous location where we propose substrates to enter the periplasmic domain in MmpL4. Common to all transport routes through the AcrAB-TolC system is the necessity of the AcrB protomers (corresponding to MmpL4 in our system) to undergo large conformational transitions resulting in the opening and closure of transport channels in the periplasmic domains to drive transport [25, 37, 38]. To achieve this, homotrimeric AcrB assumes an asymmetric conformation with pronounced structural deviations between the protomers, that reflect different protonation states of the key residues in the TMD responsible for coupling proton translocation to active transport [27, 37]. However, apart from the tilted CCDs, the MmpL4 trimer in the multimeric structure presented in this work is highly symmetric (Figure 3). Likewise, no notable asymmetries were reported for the MmpL5 and the MSEMG_1382 trimers [16, 20, 21]. Instead, we noted pronounced conformational differences of MmpL4 in its monomeric versus its trimeric context which are accompanied by conformational changes to the proton-translocating DY-pair residues (Figure 4). In addition, the MmpL3 structures feature a periplasmic cavity open towards the periplasm and the outer membrane (Figure 4).

Taking structural observations on MmpL4, MmpL5 and MmpL3 together and inferring from the well-studied RND transporter AcrB, we propose an aggregated, speculative transport mechanism for MmpS4-MmpL4 (Figure 7). Multimerization of MmpL4 with MmpS4 forms a substrate binding cavity extending from a groove formed by TM1 and TM2 in MmpL4 (Step 1 in Figure 7). In its tilted form, the CCD is unlikely transport-competent, hence it first has to reach its upright conformation (Step 2 in Figure 7), a process that is supported by MmpL4’s β-sandwich domain. It is likely that bedaquiline enters the periplasmic cavity from the outer leaflet of the cytosolic membrane. In contrast, mycobactin has to first cross the cytosolic membrane from its cytosolic binding pocket (experimentally identified in cryo-EM structures of the MmpL4 monomer [15]) to reach the periplasmic binding site (Steps 3 and 4 in Figure 7). Once the substrate is accomodated in the periplasmic binding pocket, we propose that protonation events at the TMDs involving the conserved DY-pairs (Figure 4C-E) lead to a conformational shift in the periplasmic domains, thereby guiding the previously bound substrates towards the CCD (Steps 5 and 6 in Figure 7). This hypothetical conformational change is mainly inferred from the peristaltic transport mechanism of AcrB [37] and the alternate cavity conformation experimentally seen in MmpL3 [4] (Figure 4E). Whether the substrates are transported through the CCD channel or along its surface potentially involving a mechanism similar to LPS transport in Gram-negative bacteria [39] is currently unclear. Finally, proton release is proposed to prepare the efflux system for another cycle (Step 7 in Figure 7).

Further studies will be required to fully uncover the conformational cycle of MmpL proteins in their multimeric assemblies, which will be crucial to understand the mechanism(s) by which these unique multi-subunit efflux systems export mycobactin as well as clinically important TB drugs.

## METHODS

### Strains, media and antibiotics

The *Escherichia coli* K-12 strain MC1061 was used for cloning and heterologous protein expression. Luria Broth (LB) liquid cultures and agar were used for cloning and Terrific broth (TB) liquid culture for protein expression. In *E. coli*, 100 μg ml^−1^ ampicillin and 200 μg ml^−1^ hygromycin were added to the medium when required. *M. tuberculosis* strains were grown at 37 °C in Middlebrook 7H9 liquid medium (Difco Laboratories) supplemented with 0.2 % glycerol, 10 % ADS, 0.2 % casamino acids, 24 μg/ml pantothenate, or 0.02 % tyloxapol and 20*u*M hemin. Hartmans de Bont (HdB) minimal medium (500 μM MgCl_2_, 7 μM CaCl_2_, 1 μM NaMoO_4_, 2 μM CoCl_2_, 6 μM MnCl_2_, 7 μM ZnSO_4_, 1 μM CuSO_4_, 15 mM (NH_4_)_2_SO_4_, 12 mM KH_2_PO_4,_ pH 6.8) containing 1 % (v/v) glycerol was supplemented with 0.2 % casamino acids, 24 μg/ml pantothenate. 0.02 % tyloxapol and 20 μM hemin and was used for the Alamar Blue assays. Hygromycin was added at a concentration of 50 µg ml^-1^ if needed.

### Model predictions using AlphaFold2 and AlphaFold3 and channel predictions

The structure of the MmpS4-MmpL4 trimeric complex was predicted using AlphaFold2 (Parafold setup) in multimer mode [23, 40]. The structures of CCDs of MmpL proteins were predicted using the web interface of AlphaFold3 [41] by predicting the respective hexameric MmpS-MmpL (in case a MmpS protein is part of the complex) or trimeric MmpL complexes. For simplicity, we only show the CCDs of these structures. The channel properties of the predicted CCD of MmpL4 were analyzed using the MOLEonline tool [29].

### Cloning of mmpS4-mmpL4 cross-linking variants

The gene for *mmpS4*-*mmpL4-cys-depleted* (wherein C39, C319, C335, C393 and C780 of MmpL4 are substituted by serines) was synthesized by Twist Biosciences into the fragment exchange (FX) cloning compatible entry vector pINIT (Addgene #46858) [42]. The last 123 nucleotides of *mmpL4* were codon optimized for synthesis due to their high GC content. The full nucleotide sequence of cys-depleted *mmpS4/mmpL4* is shown in Table 2. The *mmpS4*-*mmpL4-cys-depleted* gene was subcloned into the expression vector pBXC3GH (Addgene #47070) by FX cloning [43]. Nine cysteine pairs were introduced into pBXC3GH_*mmpS4-mmpL4_cys-depleted* following the QuikChange site-directed mutagenesis protocol using the primers listed in Table 3. For example, for the W6C^MmpS4^-P935C^MmpL4^ cysteine pair first the W6C^MmpS4^ mutation was introduced to *mmpS4* using the primers mmpS4_W6_Fw and mmpS4_W6_Rv. After confirmation by Sanger sequencing, the P935C^MmpL4^ mutation was introduced into *mmpL4* using the primers mmpL4_P935_Fw and mmpL_P935_Rv. The final constructs were validated by Oxford Nanopore sequencing. A 3xFLAG tag was introduced to the pML4324 vector by Gibson assembly using primers FLAG_fw and FLAG_rv for detection of protein production levels by Western Blotting. Then the *mmpS4/mmpL4-cys-depleted* was subcloned into the pML4324_3xFLAG vector by Gibson assembly. Functionally analyzed cysteine mutants were introduced directly into this complementation vector using QuikChange site-directed mutagenesis.

**Table 2:**
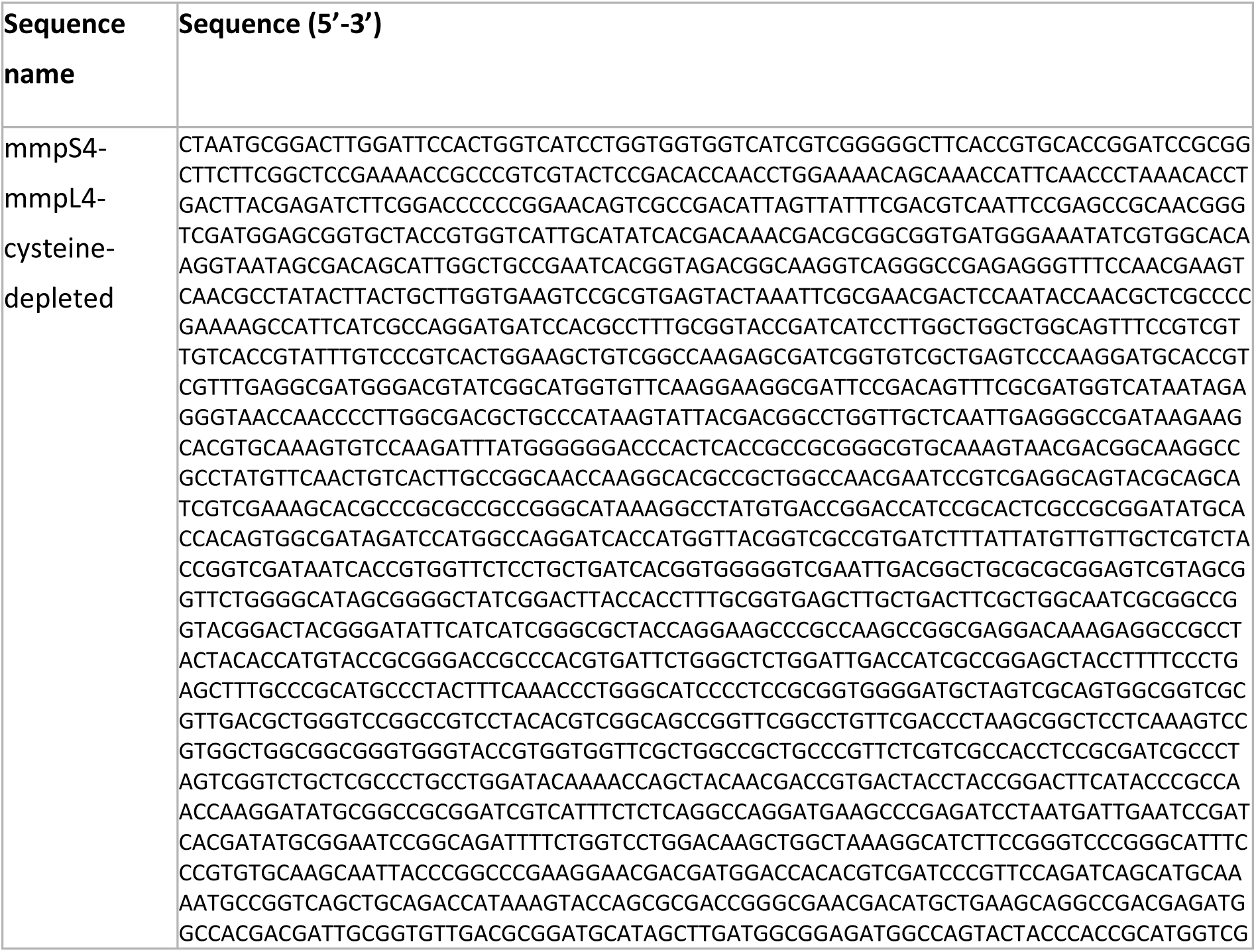
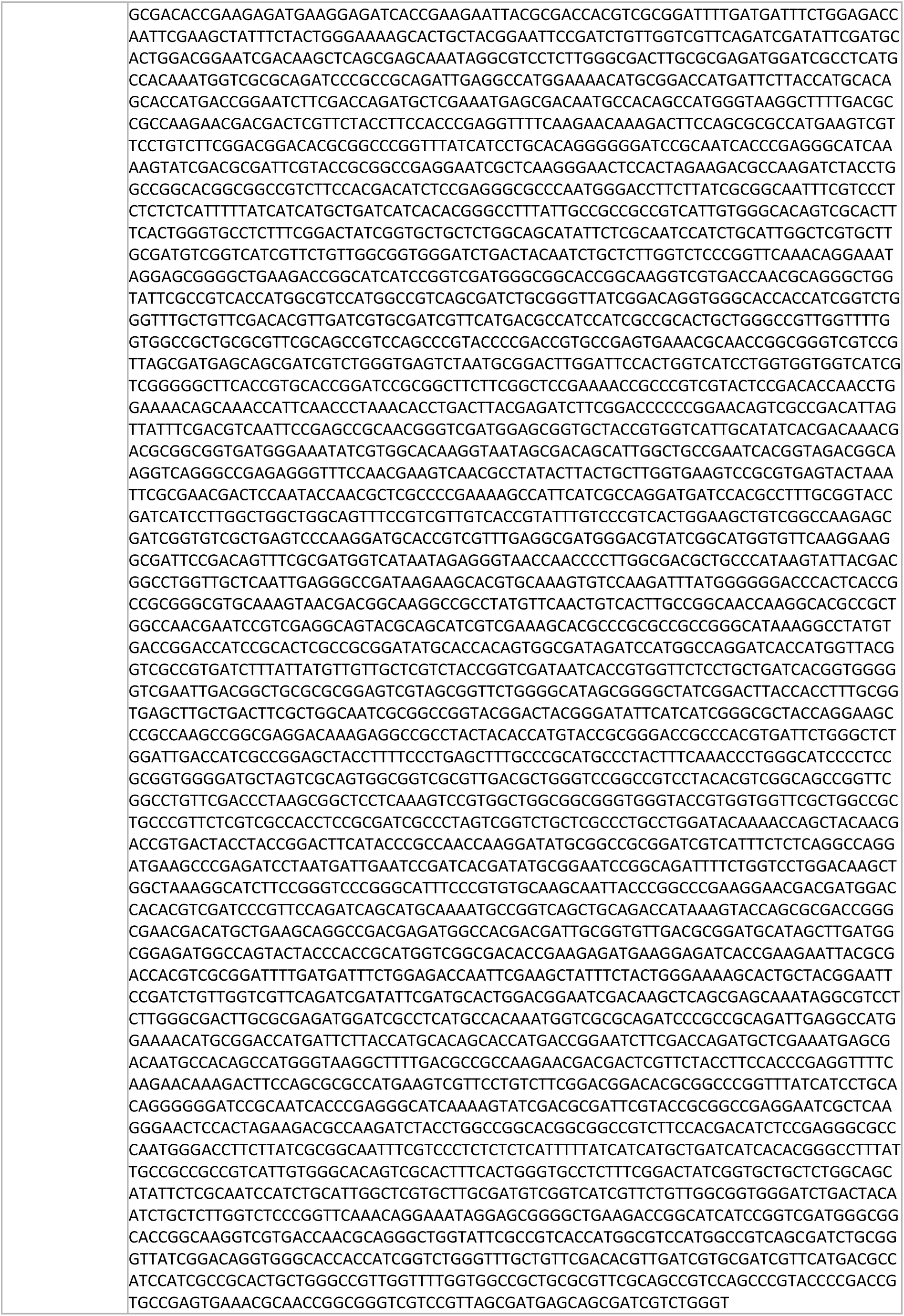
Nucleotide sequence of *mmpS4-mmpL4_cys-depleted*.

**Table 3:**
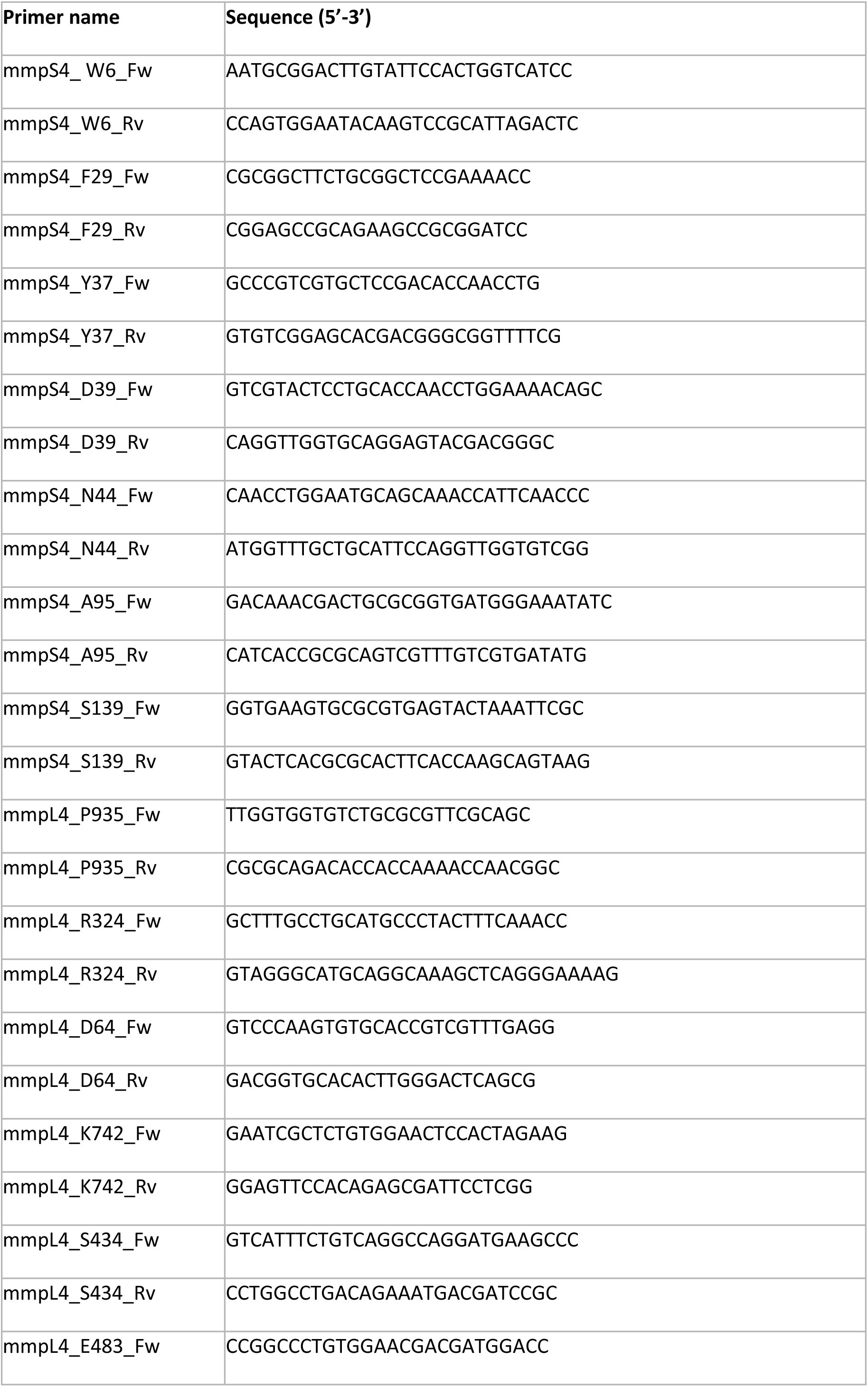
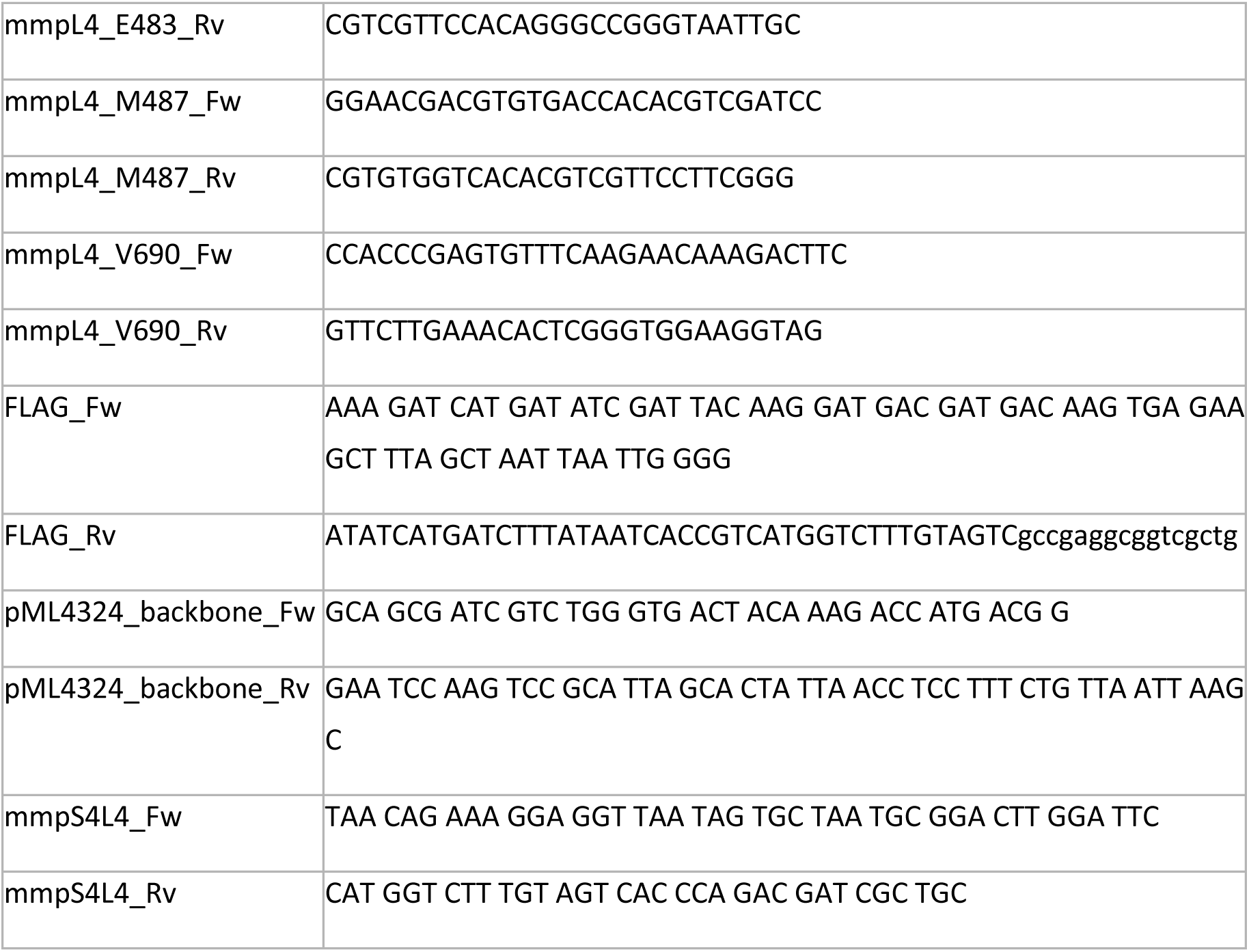
Primers used in this study.

### Cross-linked MmpS4-MmpL4 purification

The MmpS4-MmpL4 cysteine cross-linking constructs were expressed in *E. coli* MC1061. Culture volumes were either 0.6 L TB supplemented with 100 µg/ml ampicillin for small scale purification to determine crosslinking efficiency or 3 L medium to purify sufficient material for cryo-EM analyses. Cells were incubated at 37 °C for 2 h and 25 °C for 40 minutes after which protein expression was induced with 0.02 % L-arabinose. Cells were harvested after 16 hours by centrifugation for 15 minutes at 4 °C and 8,000 g in a F9-6x1000 LEX centrifuge rotor (ThermoScientific) and resuspended in PBS, pH 7.2. DNase I was added and cell were lysed by three to four passages through a Microfluidizer M-110P (Microfluidics) at 25 kPa on ice. Cell debris was removed by centrifugation for 15 minutes at 4 °C and 10,000 g in a Sorvall SLA-1500 rotor (ThermoScientific). Membrane vesicles were harvested by ultracentrifugation for 2 h at 170,000 g in a Beckman Coulter ultracentrifuge using a Beckman Ti45 rotor and resuspended in 10 or 30 ml PBS, pH 7.2 supplemented with 10 % glycerol. Prior to solubilization, 10 mM N-ethyl-maleimide was added and the resuspended membranes were rotated at 22 °C for 2 h. Then proteins were extracted by solubilizing the membranes using 1 % (w/v) β-DDM (Glycon Biochemicals) for 2 h at 4 °C. Unextracted material was harvested by ultracentrifugation at 170,000 g for 20 min and 0.5 ml of NHS activated Sepharose 4 Fast Flow (Cytiva) coupled to the anti-GFP nanobody (PDB-ID: 3K1K) was added to the supernatant to capture the GFP tagged MmpL4-MmpS4 complex. All purification steps were carried out at 4 °C. Batch binding was performed for 2 h, after which the resin was washed with 20 ml of PBS, 10 mM NEM, 10 % glycerol, 0.03 % β-DDM followed by 50 ml TBS (20 mM Tris/HCl, pH 7.5, 150 mM NaCl), 10 % glycerol, 0.03 % β-DDM. The resin was incubated with 100 µg 3C protease for 2 h to elute the protein by cleaving off the GFP-His_10_-tag. The flow through was collected and the resin was washed with 2-4 ml buffer to collect the cleaved protein. The sample was analyzed by reducing and non-reducing SDS-PAGE gel electrophoresis to assess crosslinking efficiency. Proteins were concentrated with 100 kDa MWCO concentrator (Merck Millipore) to 1-2 mg/ml and flash frozen and stored at -80 °C or immediately purified by size-exclusion chromatography on a Superose 6 Increase 10/300 GL column (Cytiva) in TBS, 0.03 % β-DDM.

### Cryo-EM sample preparation and data acquisition

The cross-linking construct #5 of the MmpS4-MmpL4 complex was concentrated to 4 mg/ml and applied to glow discharged (0.39 mbar, 15 mA, 1 min) holey carbon grids (QuantiFoil R1.2/1.3 Au 300 mesh). Excess sample was removed by blotting at 4 °C for 4.5-5.5 s with a relative humidity of 100 % and a blotting force of 20 using a Vitrobot Mark IV (ThermoFisher). The sample was vitrified by plunging into ethane/propane. Cross-linked MmpS4-MmpL4 was imaged on a Titan Krios G3i (300 kV, 100 μm objective aperture) equipped with a BioQuantum energy filter (20 eV energy filter slit) and a K3 direct electron detector (6k x 4k pixels). EPU 2.9 was used for automated data acquisition of dose-fractionated micrographs in a defocus range of -1 – -2.2 μm in 0.2 µm increments. Micrographs were recorded at a magnification of 130,000x, with a pixel size of 0.65 Å and 1.3 s exposure time in counted super-resolution mode and Fourier cropped by 2. A total of 14588 micrographs were recorded for the cross-linked MmpL4-S4 complex #5 with a dose of 21.10 e^−^/px/sec resulting in a total electron exposure of 62.92 e^−^/Å^2^.

### Cryo-EM image processing, model building and refinement

The acquired data was processed in CryoSPARC4.7.0 [44]. Data quality was monitored and initial processing steps including patch motion correction and patch CTF estimation were performed in CryoSPARC live. Good micrographs were selected based on the resolution of the estimated CTF fit and relative ice thickness. Particles were selected and filtered in two different ways (Figure 3 — figure supplement 2). First, particles were picked using the blob picker, extracted with a box size of 400 pixel and Fourier cropped to 100 pixels (2.6 Å per pixel). Two rounds of 2D-classification were performed, followed by one ab-initio reconstruction and one additional round of 2D-classification. In a parallel approach, micrographs were first denoised and then particles were picked using the blob picker. The resulting particles were selected in the same way as described above, with two rounds of 2D-classification, one round of ab-initio reconstruction and one additional round of 2D-classification. The final selection of particles from both approaches were merged and duplicates were removed. Particles were unbinned, reextracted, and an ab-initio reconstruction was performed. The subsequent non-uniform (NU) refinement [45], was performed once with C1 symmetry and once with C3 symmetry constraints. NU refinement using C1 symmetry resulted in a 3.09 Å map of the MmpL4-S4 complex with a “short” CCD. NU refinement using C3 symmetry resulted in a 2.89 Å map of the MmpL4-S4 complex. The C1-symmetrical map was locally refined with a mask created from its CCD. The local refinement resulted in a 3.22 Å map of the MmpL4-S4 complex with a “long” CCD. Resolutions are stated according to the 0.143 cut-off criterion. Local resolution estimation was carried out in CryoSPARC.

### Model building

The monomeric MmpL4 model (PDB: 9GI0) was docked into the C3 symmetrical density obtained for the MmpL4-S4 complex using PHENIX [46], Coot [47] and ISOLDE [48]. This model was used as input for the C1-symmetrical model with “short” and “long” CCD and the AlphaFold3 prediction of the CCD of the MmpL4-S4 complex was docked into the tilted CCD density. Cavities of MmpL3, the monomeric MmpL4, and the MmpL4-S4 complex were calculated using a local installation of the 3 V package (shell probe radius: 5 Å, solvent-excluded probe radius of 1.4 Å, grid spacing 1.3) [26].

### Mycobactin toxicity susceptibility assay

The susceptibility of the parent *Mtb* strain mc^2^6230 [49], the *Mtb* Δ*mmpL4/mmpL5* strain and the *Mtb ΔmmpL4/mmpL5* strains complemented with the wt and mutated *mmpS4-mmpL4 operons* to mycobactin was measured using a microplate Alamar Blue assay (MABA) as described previously with modifications [9]. Cultures of the *Mtb*Δ*mmpL4/mmpL5* strain and the complemented strains were grown in supplemented 7H9 medium to an OD_600_ of 1. The cultures were then filtered using a 5 µM filter to remove clumps. Then, the cells were centrifuged and the cell pellet was washed with Hartmans de Bont (HdB) medium and centrifuged again. The pellet was resuspended in HdB medium supplemented with 0.2 % casamino acids, 24 μg/ml pantothenate, 20 μM hemin and 0.02 % tyloxapol. The cell suspension was adjusted to an OD_600_ of 0.05 to obtain a final starting OD_600_ of 0.025 in the culture wells by mixing 100 µl of the cell suspension with 100 µl of mycobactin-containing medium. The cells were then incubated in microplates with increasing mycobactin concentrations for seven days at 37 °C. Resazurin (90 μM) was added and cells were incubated for an additional six hours after which the fluorescence of the metabolically converted resazurin dye was measured as an indicator of cell viability.

### Western blotting

Complemented *Mtb* Δ*mmpL4/mmpL5* strains were grown to an OD_600_ of ∼2 in Middlebrook 7H9 medium (Difco Laboratories) supplemented with 0.2 % glycerol, 10 % ADS, 0.2 % casamino acids, 24 μg/ml pantothenate, 0.02 % Tyloxapol supplemented with 20 μM hemin. Cultures were harvested and washed twice with PBS and then resuspended in PBS. Bead beating was carried out using a Fast Prep FP120 Homogenizer with intermittent incubations on ice. After the first cycle of bead beating, 1 % SDS was added to the cells followed by two cycles of bead beating. Cell lysates were subsequently centrifuged at 3200 x g to remove cell debris, resulting in the isolation of the whole cell lysate (WCL). The protein content in the lysates was determined using the Bradford (Bio-Rad) assay. For gel electrophoresis 5 μg of WCL were mixed with protein loading dye (either with or without β-mercaptoethanol) and separated on a 10 % sodium dodecyl sulfate (SDS)-polyacrylamide gel. The proteins were then transferred to a polyvinylidene difluoride membrane using the Trans-Blot Turbo Transfer System (Bio-Rad). The blots were blocked with 5 % skim milk (BD) in TBS containing 1 % Tween-20 (TBS-T) overnight and incubated with the rabbit polyclonal anti-flag (1:5000, Sigma-F7425, 069K4767). After three washes with TBST, the blots were incubated with the goat anti-rabbit IgG-HRP (1:10000, Sigma) antibody. All antibody dilutions were done in TBS-T. The Western blots were imaged using the Li-Cor Odyssey or Gel Doc XR+ (Bio-Rad).

## DATA AVAILABILITY

Structural models have been deposited to the Protein Data Bank (PDB) with the accession codes 9SYJ (data processed using C1 symmetry), 9SYT (data processed using C1 symmetry and applying a mask to improve the resolution of the CCD), and 9SYV (data processed using C3 symmetry). The cryo-EM data has been deposited in the Electron Microscopy Data Bank (EMDB) with the accession codes D_1292151303, D_1292151414, and D_1292151419 respectively. Plasmids and other data that support the findings of this study are available from the corresponding authors upon request.

## ACKNOWLEDGMENTS

Work in the laboratory of M.A.S. was funded by the European Research Council (ERC) (consolidator grant n° 772190) and the Swiss National Science Foundation (PP00P3_170625). Work in the laboratory of M.N. was supported by the National Institutes of Health grant R01 AI137338. A.A.G. received support from the Forschungskredit of the University of Zurich (FK-21-041). We thank Charlott Stock for performing the AlphaFold2 runs on the EMCC cluster https://emcc.au.dk, and Martina Peter for assistance in the cryo-EM data collection. Imaging was performed with equipment maintained and with the support by the Center for Microscopy and Image Analysis, University of Zurich. We gratefully acknowledge EMCC and Aarhus University for providing computational resources to run AlphaFold2 predictions.

## AUTHOR CONTRIBUTIONS

J.C.E., M.N., and M.A.S. conceived the project. J.C.E. generated the cysteine variants, performed the cross-linking analysis, purified the cross-linked complex #5, and prepared cryo-EM grids. N.P.L. measured cryo-EM data, processed the data with help of A.A.G., and built the model with support of A.A.G.. V.M. performed mycobactin toxicity assays and Western blot analyses. J.C.E., N.P.L., M.N., and M.A.S. interpreted the data. N.P.L. prepared the figures with inputs from J.C.E., and M.A.S.. N.P.L., J.C.E, M.N., and M.A.S. wrote the manuscript with input from all other authors.

## SUPPLEMENTARY INFORMATION

**Figure 1 – figure supplement 1.**
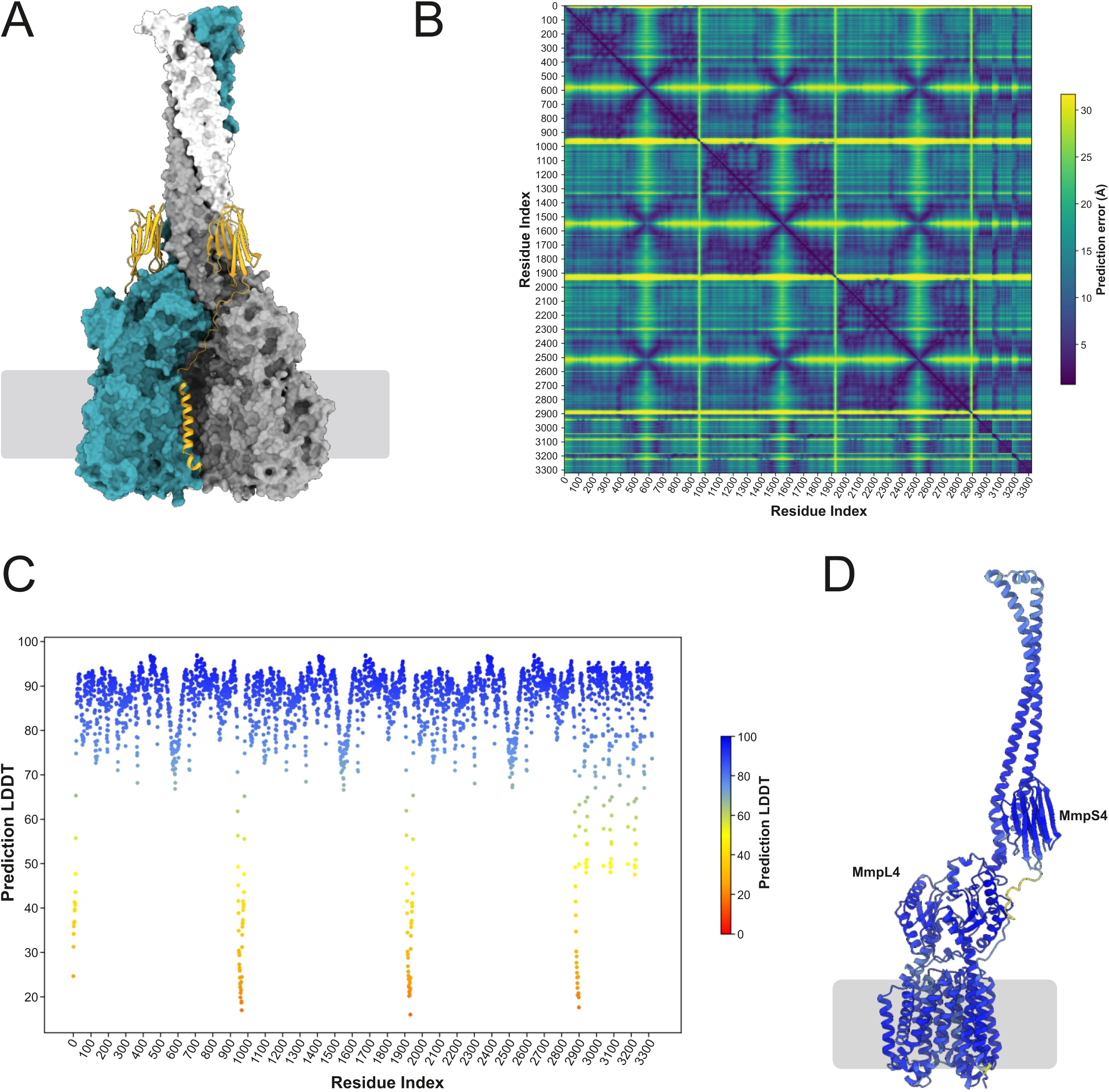
AlphaFold2 prediction of the hexameric (MmpS4)_3_-(MmpL4)_3_ complex. (**A**) The three protomers of MmpL4 are shown in surface presentation and colored teal, grey and white. The MmpS4 protomers are shown as yellow cartoon (only two of them are visible). The boundary of the cytoplasmic membrane is indicated by grey lines. (**B**) Predicted alignment error (PAE) plot of the hexameric (MmpS4)_3_-(MmpL4)_3_ complex reported by AlphaFold2. It shows the expected distance error (in Å) between each residue in the predicted model. The three MmpL4 protomers cover residues 1-2901, and the three MmpS4 protomers residues 2902-3321. (**C**) Predicted Local Distance Difference Test (pLDDT) plot. The three MmpL4 protomers cover residues 1-2901, and the three MmpS4 protomers residues 2902-3321. (**D**) Depiction of pLDDT values in the context of the predicted MmpL4/MmpS4 structure (only first protomer shown, N- and C-terminus of MmpL4 cleaved).

**Figure 2 – figure supplement 1.**
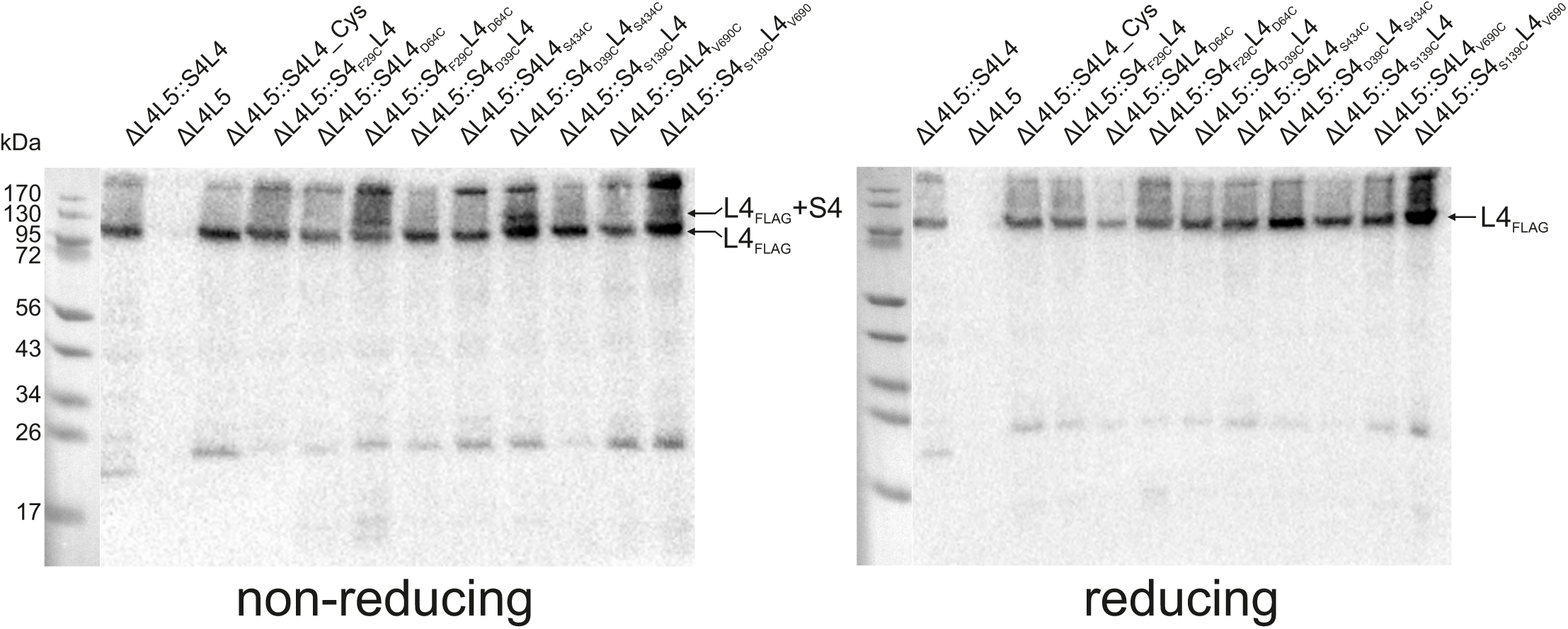
Analysis of crosslink formation between MmpL4 and MmpS4 cysteine mutants in *M. tuberculosis*. Western blot analysis of cell lysates from the indicated *Mtb* strains using an anti-FLAG_3_ antibody (see methods) treated with (reducing) or without (non-reducing) β-mercaptoethanol. The bands corresponding to FLAG-tagged MmpL4 (L4_FLAG_) and FLAG-tagged MmpL4 crosslinked to MmpS4 (L4_FLAG_+S4) are indicated with an arrow.

**Figure 3 – figure supplement 1.**
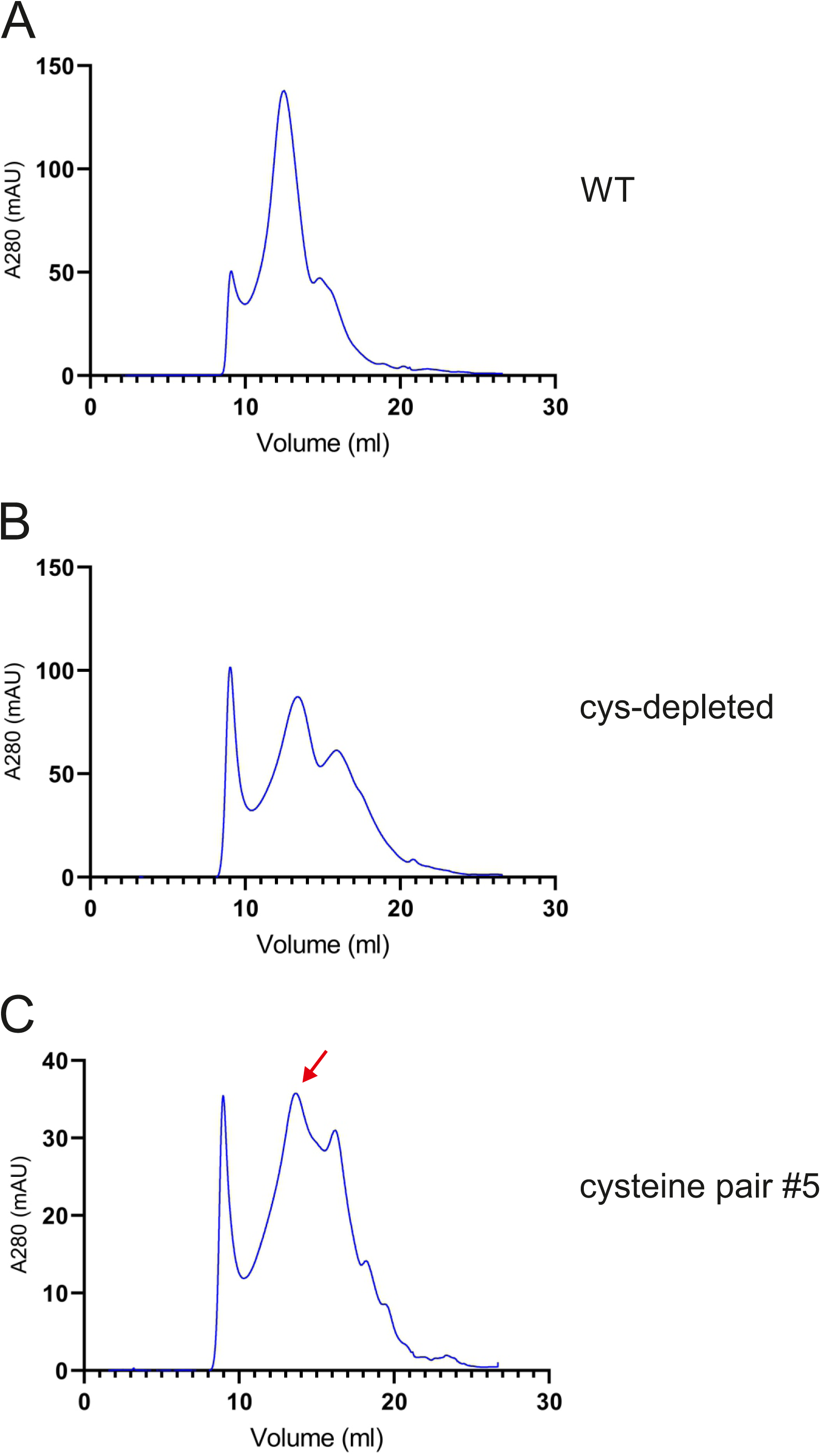
Size exclusion chromatograms. (**A**-**C**) Size exclusion chromatography traces of wild-type MmpS4-MmpL4 (**A**), cys-depleted MmpS4-MmpL4 (**B**) and MmpS4-MmpL4 containing the cysteine pair #5 (**C**). The proteins were separated on a Superose 6 Increase 10/300 GL column. The main peak of MmpS4-MmpL4 containing the cysteine pair #5 eluting at 13.6 ml (indicated by red arrow) was concentrated and analyzed by cryo-EM. The cryo-EM structure of the hexameric (MmpS4)_3_-(MmpL4)_3_ complex was determined from this sample.

**Figure 3 – figure supplement 2.**
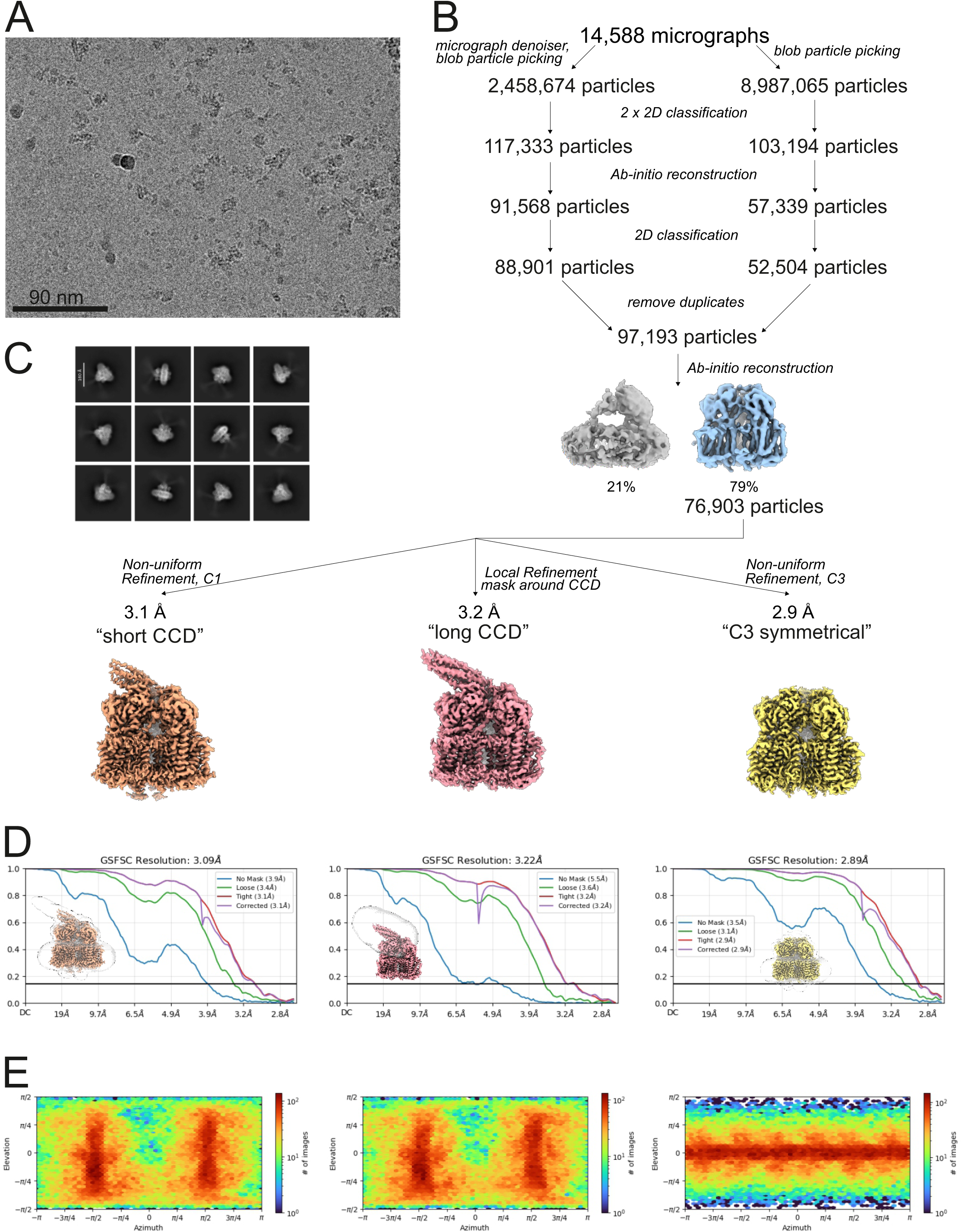
Cryo-EM reconstruction of the hexameric MmpS4-MmpL4 complex. (**A**) Representative cryo-EM micrograph. (**B**) Image processing work flow. (**C**) 2D-class averages. (**D**) FSC plot used for resolution estimation. (**E**) Angular distribution plot of particles included in the final 3D reconstruction.

**Figure 3 – figure supplement 3.**
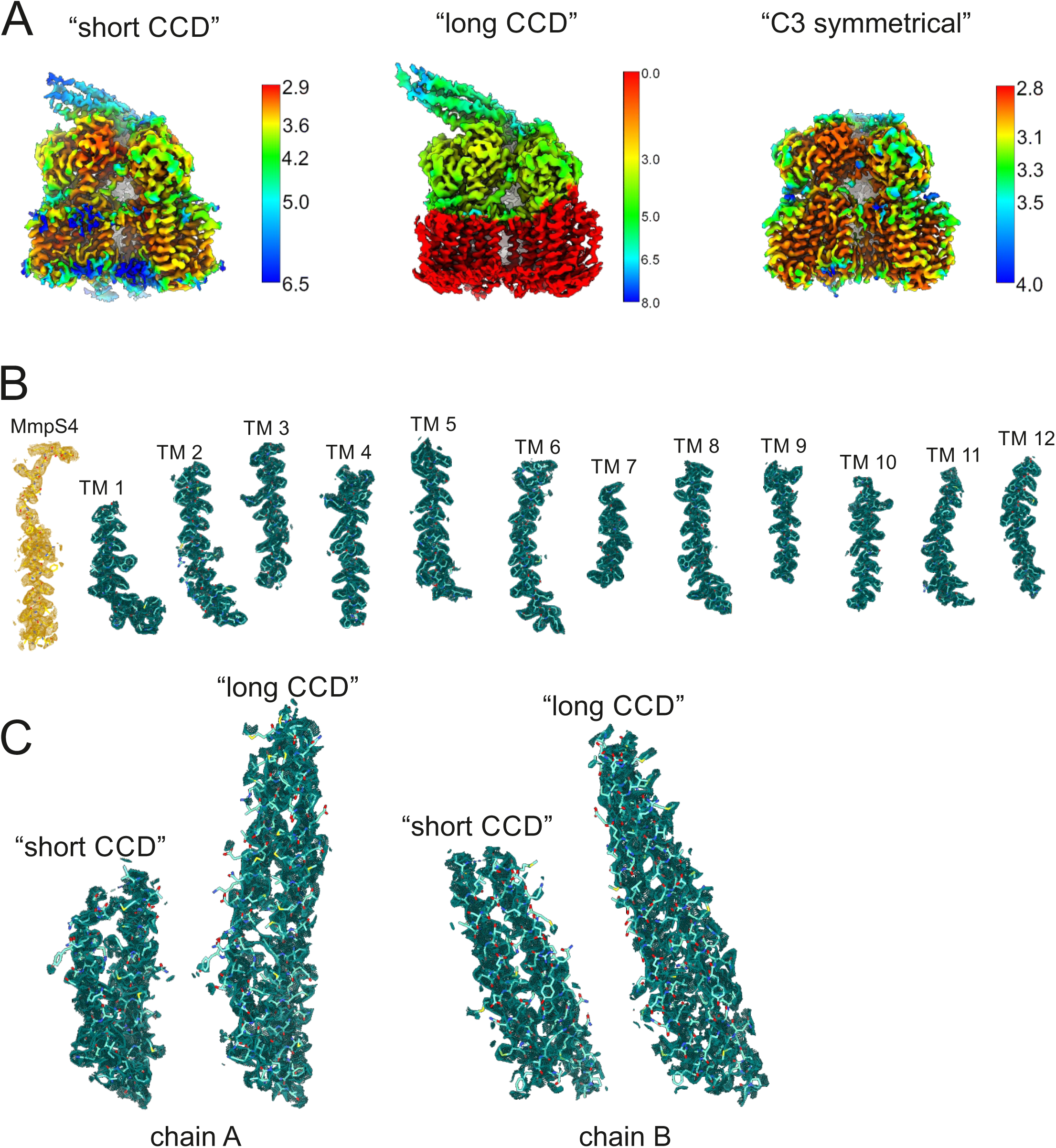
Local resolution estimations and cryo-EM densities of the hexameric MmpS4-MmpL4 complex. (**A**) Final reconstructed maps colored by local resolution. Note that for the “long CCD” map, a large part of the map was not part of the mask and thus no local resolution could be determined (red parts, resolution arbitrarily set to zero). (**B**) Cryo-EM densities of the hexameric MmpS4-MmpL4 complex “short CCD” with the respective refined model superimposed. The model is shown as sticks and structural elements are labelled. Transmembrane helices (TM) are colored in green, MmpS4 in yellow. Densities were contoured at 5.4σ. (**C**) Cryo-EM densities of CCDs from “short CCD” and “long CCD” models, chains A on the left and chains B on the right. Densities were contoured at 4.2σ. Cryo-EM reconstruction of the hexameric MmpS4-MmpL4 complex.

**Figure 3 – figure supplement 4.**
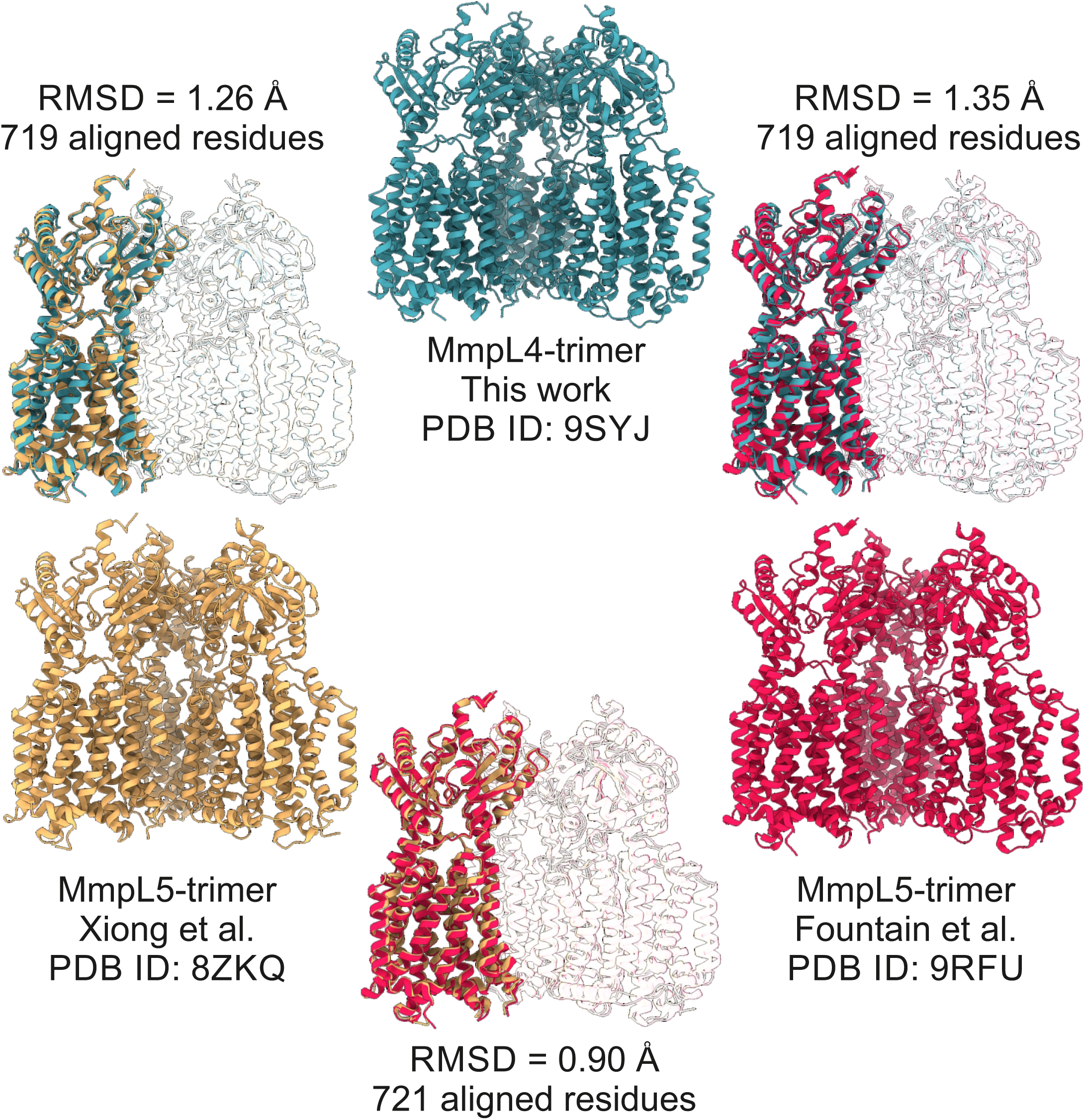
Structural comparisons of trimeric *M. tuberculosis* MmpL4 and MmpL5. The first protomers of the trimeric MmpL4 structure (this study, PDB ID: 9SYJ, chain A) and the two trimeric MmpL5 structures (Xiong et al., PDB ID: 8ZKQ, chain A / Fountain et al., PDB ID: 9RFU, chain D) were superimposed using the matchmaker tool of ChimeraX (Needleman-Wunsch alignment algorithm, BLOSUM-62 similarity matrix). The respective RMDS values are depicted directly in the figure, along with the number of aligned residues.

**Figure 4 – figure supplement 1.**
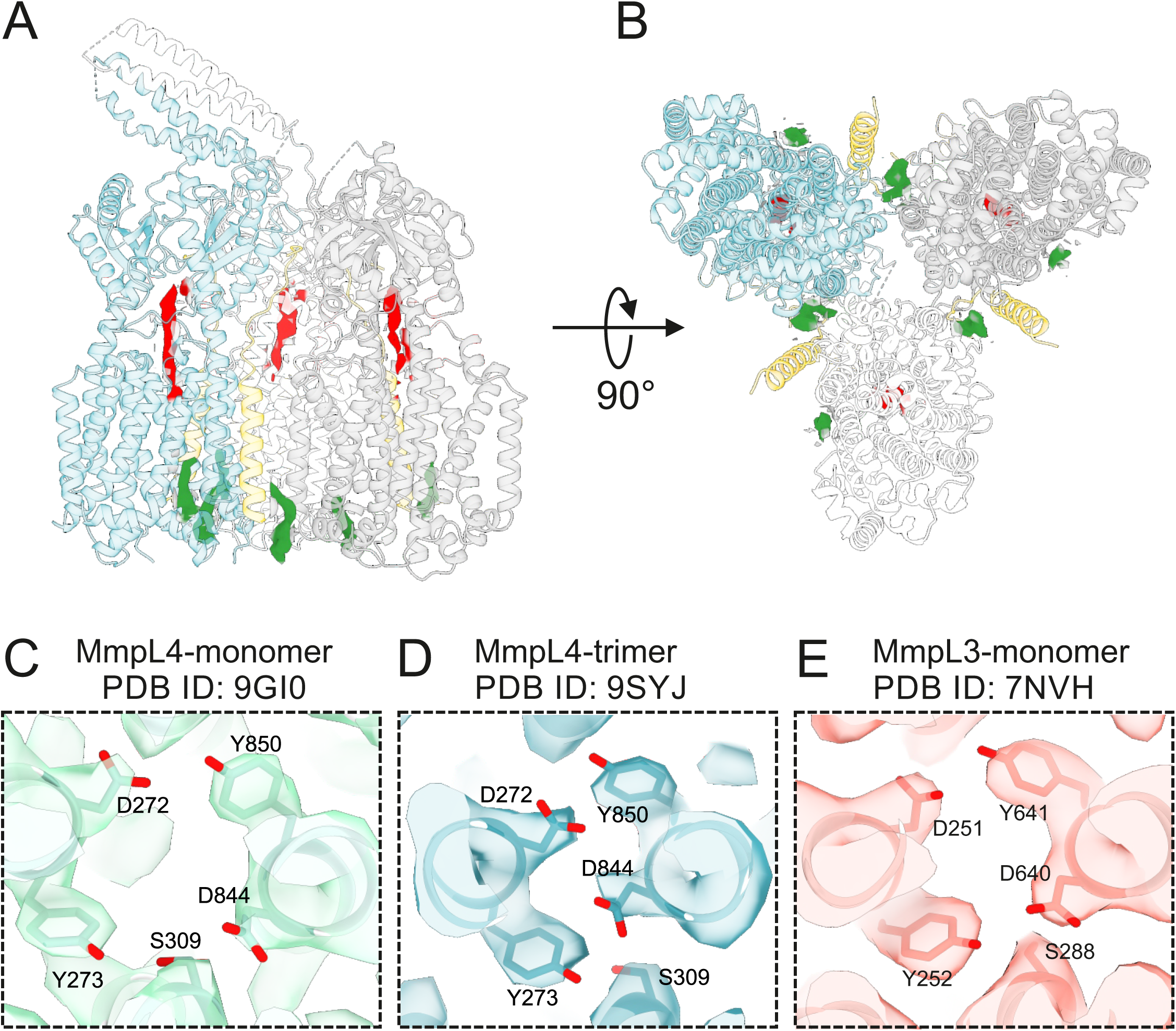
Analysis of non-proteinaceous densities and cryo-EM densities of DY-pairs. (**A** and **B**) Systematic analysis of cryo-EM densities (contoured at 7.4σ) not accounted by the MmpS4-MmpL4 polypeptide chains in the context of the asymmetric structure processed in C1 shown from the side (**A**) and from the cytosol (**B**). Non-proteinaceous densities in the periplasmic domains marked in red correspond to bound β-DDM. Densities at the level of the cytosolic leaflet of the membrane marked in green most likely correspond to partially ordered lipids of unknown identity. (**C** – **E**) The conserved DY-pairs and S309(MmpL4)/S288(MmpL3) are shown as sticks and viewed from the periplasm as in main Fig. 3. (**C**) Monomeric MmpL4 (PDB ID: 9GI0) contoured at 11.9σ. (**D**) Trimeric MmpL4 (this work, PDB ID: 9SYJ, chain A) contoured at 8.9σ. (**E**) Monomeric MmpL3 (PDB ID: 7NVH) contoured at 4.5σ.

**Figure 6 – figure supplement 1.**
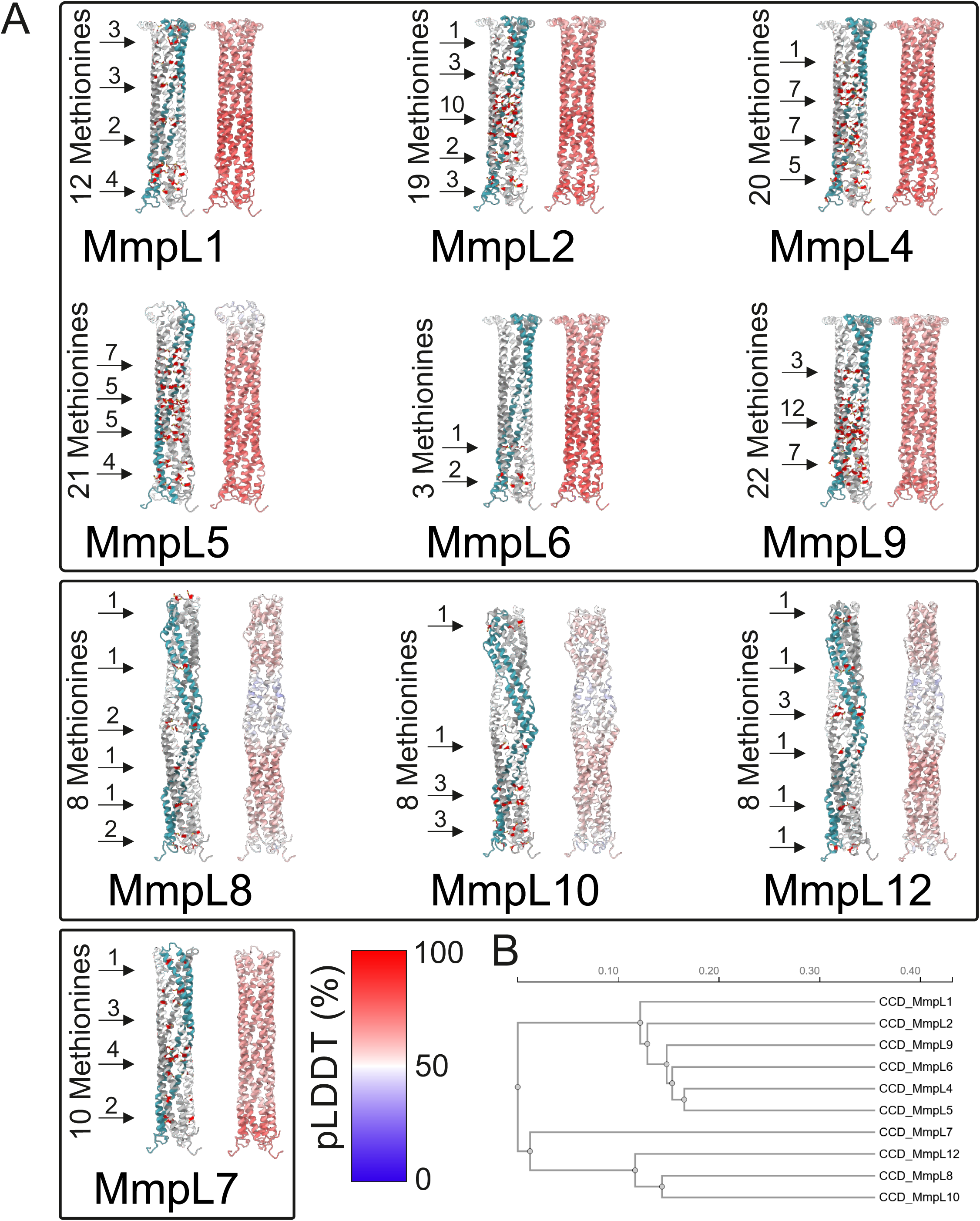
Comparative analysis of the coiled-coil domains of MmpL proteins of *M. tuberculosis*. (**A**) CCDs of *M. tuberculosis* MmpL proteins shown as cartoons. The models are AlphaFold3 predictions of the respective MmpS-MmpL hexamers (in case an MmpS protein is present) or the MmpL trimers. For each CCD, methionine residues are shown as red sticks and the number of methionine residues per α-helical hairpin is provided in the pictures on the left and in the pictures on the right, the models are colored according to their Predicted Local Distance Difference Test (pLDDT) values. The coloring scheme is provided at the bottom. (**B**) Phylogenetic tree of the CCDs shown in (**A**).

